# Mechanical Stretch Kills Transformed Cancer Cells

**DOI:** 10.1101/491746

**Authors:** Ajay Tijore, Mingxi Yao, Yu-Hsiu Wang, Yasaman Nematbakhsh, Anushya Hariharan, Chwee Teck Lim, Michael Sheetz

## Abstract

Transformed cancer cells differ from normal cells in several important features like anchorage independence, Warburg effect and mechanosensing. Consequently, transformed cancer cells develop an anaplastic morphology and respond aberrantly to external mechanical forces. Consistent with altered mechano-responsiveness, here we show that transformed cancer cells from many different tissues have reduced growth and become apoptotic upon cyclic stretch as do normal cells after the transformation. When matrix rigidity sensing is restored in transformed cancer cells, they survive and grow faster on soft surface upon cyclic stretch like normal cells but undergo anoikis without stretch by activation of death associated protein kinase1 (DAPK1). In contrast, stretch-dependent apoptosis (mechanoptosis) of transformed cells is driven by stretch-mediated calcium influx and calcium-dependent calpain 2 protease activation on both collagen and fibronectin matrices. Further, mechanosensitive calcium channel, Piezo1 is needed for mechanoptosis. Thus, cyclic stretching of transformed cells from different tissues activates apoptosis, whereas similar stretching of normal cells stimulates growth.

## Introduction

The transformed phenotype was initially described in early studies of cancer cell growth, since tumor cells would often grow on soft agar plates, while normal cells from the same tissue required a rigid surface for growth (Hamburger and Salmon, 1977). Recent studies have shown that transformed cancer cells from many different tissues lack rigidity sensors. Restoration of rigidity sensing, through cytoskeletal protein expression in cancer cells, blocks transformed growth (Wolfenson et al., 2016; Yang et al., 2018). Further, normal cells can become transformed by depleting cytoskeletal proteins that are required for rigidity-sensing. This raises many questions regarding the transformed phenotype, since altering cytoskeletal protein levels in cancer cell lines restores rigidity-dependent growth and reciprocal alterations in normal cells cause transformed growth in many different tissue backgrounds. As the morphology and mechanical properties of the transformed cancer cells are dramatically different from the normal cells, there may be substantial differences in the mechano-sensitivity between normal cells and transformed cancer cells.

Several reports have indicated that cancer cell growth is vulnerable to mechanical forces. Studies showing inhibition of tumor growth after stretching or exercise in mice model could be explained through a mechanical force-dependent growth inhibition (Berrueta et al., 2018; Betof et al., 2015). Fluid shear-induced killing of circulating tumor cells and adhesive cancer cells can also be explained by increased sensitivity to mechanical forces (Lien et al., 2013; Regmi et al., 2017). In addition, ultrasonic and shock-wave therapies have demonstrated that cancer cells are particularly sensitive, although concerns were raised about increased metastasis and healthy tissue damage (Lin et al., 2012; Marano et al., 2017; Nicolai et al., 1994; Zequi et al., 2018). We designed this study to investigate the effect of mechanical cyclic stretch on the behavior of normal versus transformed cancer cells from a variety of different tissue backgrounds.

Because the presence or absence of a rigidity-sensing complex correlates strongly with the normal (rigidity-dependent) and transformed growth (rigidity-independent) states, respectively, it is relevant to understand the important elements of the rigidity sensors. They are transient sensors that involve the assembly of sarcomeric units of ~2 μm in length with anti-parallel actin filaments anchored to the matrix adhesion sites. Myosin IIA contraction pulls the adhesions to a constant displacement of about 100 nm for 30 seconds irrespective of substrate rigidity (Wolfenson et al., 2016). If the matrix is rigid, then the contractile force will exceed ~25 pN and the adhesions will be reinforced. Adhesion disassembly and eventual cell apoptosis will occur if the surface is soft. Many different tumor suppressors are part of the rigidity sensors including TPM2.1 (formerly known as Tm1), myosin IIA, DAPK1, α-actinin 4 and receptor tyrosine kinases like AXL and ROR2 (Helfman et al., 2008; Qin et al., 2018; Yang et al., 2016; Yang et al., 2018). A common mechanism of the malignant transformation is the depletion of TPM2.1 or the increased expression of Tpm3 (Raval et al., 2003; Stehn et al., 2013). Decreasing the ratio of TPM2.1 to Tpm3 causes the loss of matrix rigidity sensing and enables transformed cell growth. This process can occur even in the normal cells (Wolfenson et al., 2016; Yang et al., 2018). In contrast, restoration of TPM2.1 level in many cancer cells re-establishes rigidity-dependent growth and inhibits transformation in those cells (Yang et al., 2018). Other mechanosensory cytoskeletal proteins (myosin IIA, α-actinin, filamin A, AXL, ROR) are part of the rigidity sensing complex. Depletion of these sensory proteins and other proteins like scaffolding proteins (caveolin1) promotes cell transformation, while their restoration in cancer cells will block transformed cell growth (Lin et al., 2015; Wolfenson et al., 2016; Yang et al., 2016; Yang et al., 2018). Thus, the transformed state of cancer cells appears in many different tissue backgrounds to result from the loss of rigidity sensing, i.e. transformed cells grow on soft surfaces because they fail to sense the surface softness.

There are a number of common features of cancer cell mechanics that differ from normal cells, including a decrease in the rigidity of the cortical cytoskeleton (Lin et al., 2015; Swaminathan et al., 2011) and increased traction forces on matrices (Kraning-Rush et al., 2012). In addition, there are multiple reports demonstrating the altered expression and function of calcium channels as well as increased calpain activity in cancer cells (Azimi et al., 2014; Storr et al., 2011). In this study, we explore various aspects of the transformed cancer cells that cause selective apoptosis of the transformed, but not of the normal cells. Surprisingly, we find that the cyclic stretching of transformed cancer cells inhibits their growth and activates apoptosis. Restoration of matrix rigidity sensing in transformed cancer cells by TPM2.1 expression results in increased growth and survival upon cyclic stretch, particularly on soft surfaces. Transformed cell death results from increased calcium entry upon stretch that activates calpain 2 mediated-mitochondrial apoptosis.

## Results

### Cancer Cells Elongate upon Cyclic Stretch: Magnitude, Frequency and Rigidity Dependence

To understand the impact of cyclic stretch on cancer cell behavior, we used the MDA-MB-231 metastatic breast cancer cell line which is known for being highly aggressive. Cyclic stretching of MDA-MB-231 cells on fibronectin-coated flat PDMS (2 MPa, rigid) for 6 hrs caused prominent cell elongation (quantified by the cell aspect ratio (AR)) (Figure 1A and S2A). In the case of MDA-MB-231 cells, the highest AR was at 5% cyclic strain (AR ~12) in contrast to 1% (AR ~3) and non-stretched cells (AR ~2). In addition, a single stretch (5% for 6 hrs) did not alter cell elongation (AR was the same as non-stretched cells). Next, to assess the effect of stretching frequency on MDA-MB-231 cell morphology, cells were subjected to different frequencies (0.1 Hz to 1 Hz) for 6 hrs. Interestingly, cells displayed frequency-dependent cell elongation with maximum elongation observed at 0.5 Hz with 5% cyclic strain (AR ~11) (Figure 1B). On the other hand, cells showed less elongation at 0.1 Hz (AR ~4) and no significant difference in the elongation for 1 Hz vs. 0.5 Hz was observed. Thus, the cancer cell elongation in response to cyclic stretch optimally at a frequency of 0.5 Hz and a strain of 5% was chosen to minimize damage to normal cells.

**Figure 1.**
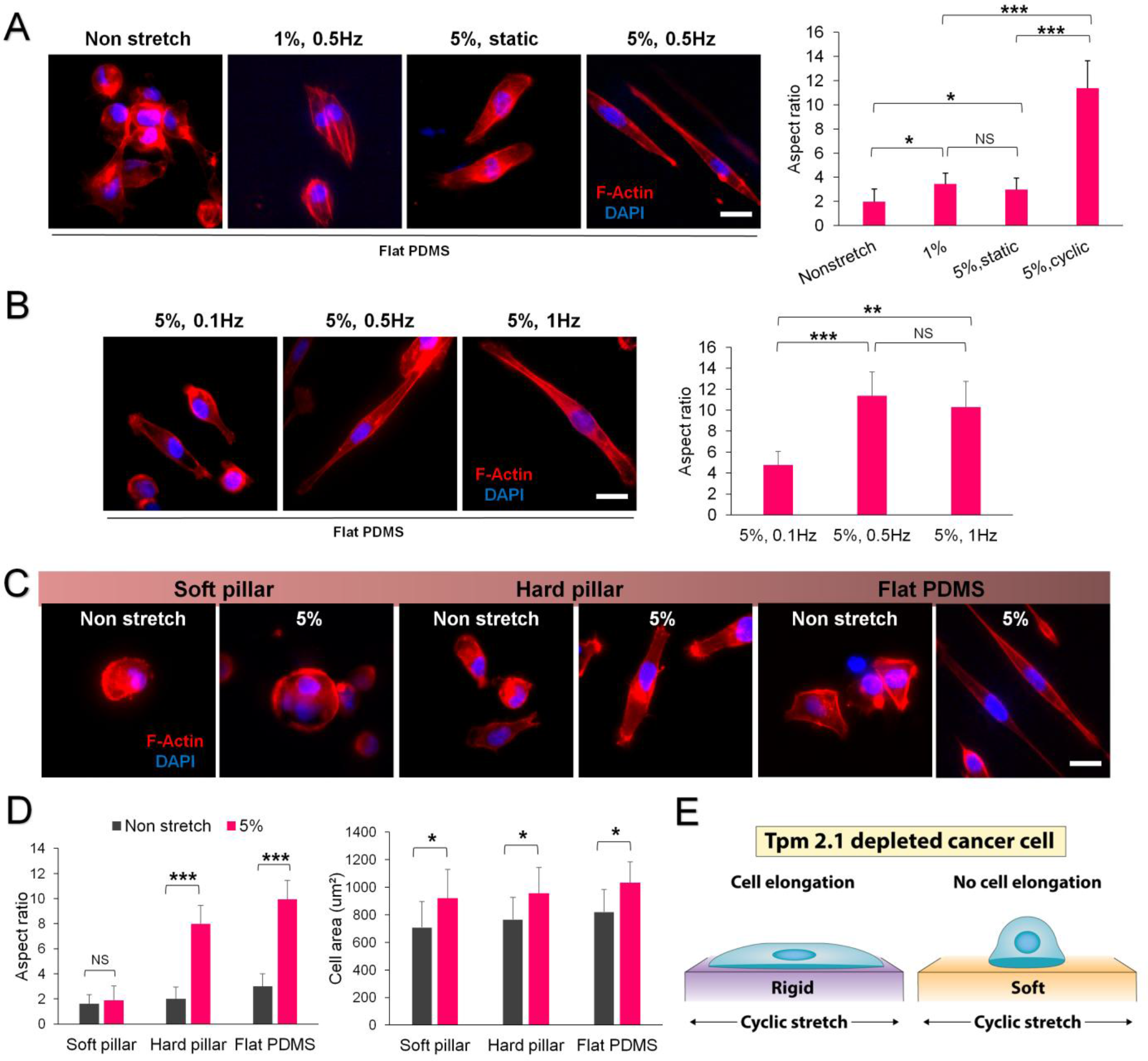
Effects of Magnitude and Frequency of Cyclic Stretching on Cancer Cell Morphology with Different Matrix Rigidities. **(A)** MDA-MB-231 cells displayed the highest elongation (AR ~12) at 5% cyclic strain on flat PDMS surfaces (rigid) after 6 hrs. **(B)** Cancer cell elongation was also found to be dependent on the frequency of stretching, demonstrating maximum elongation at 0.5Hz with 5% cyclic strain (AR ~11). **(C)** Matrix rigidity influenced the cancer cell elongation. Representative images illustrating cyclic stretch-dependent cancer cell elongation on rigid surfaces (rigid pillars and flat PDMS) but not on the soft pillar surfaces. **(D)** Statistical data showing marked increases in the cell AR on rigid surfaces only. However, a stretch-dependent cell area increase was observed on both rigid and soft surfaces after cyclic stretch. **(E)** Schematic diagram illustrating matrix rigidity-dependent cancer cell elongation upon cyclic stretch. n=100 cells for cell aspect ratio and n=30 cells for cell area. Experiments were repeated at least three times. \**P*< 0.05, \*\**P*< 0.01, \*\*\**P*< 0.001. Scale bar: 20 μm.

To investigate the effect of matrix rigidity on stretch-dependent cancer cell elongation, MDA-MB-231 cells were cyclically stretched (5%, 0.5 Hz) on surfaces with different rigidity (Figure 1C). On soft pillars (~8 kPa), cells failed to display elongation in response to cyclic stretch and showed similar AR values on stretch and non stretch surfaces (Figure 1D). In fact, cells were round on soft surfaces which resembled the morphology of non-stretched cells on soft surfaces. In contrast on rigid pillars (~55 kPa) and flat PDMS surfaces (2 MPa), cells had an elongated morphology, while non-stretched cells had a non-polarized morphology on both surfaces. In addition, we observed that cyclic stretching was responsible for a small increase in the cell area irrespective of the matrix rigidity (Figure 1D).

To delineate the influence of extracellular matrix protein (ECM) on stretch-dependent cancer cell morphology, similar stretching experiments were repeated with MDA-MB-231 cells cultured on collagen-I coated surfaces (Figure S3A). Again, cells elongated after cyclic stretch on flat PDMS surfaces (AR ~9) in comparison to non-stretched cells (AR ~3). In contrast, stretching did not result in the cell elongation on soft surfaces (AR ~3). Further, the stretch-dependent cell area increment was noticed on rigid surfaces only. In short, MDA-MB-231 cells responded similarly on fibronectin and collagen-I coated surfaces by showing elongated and round morphology on rigid and soft surfaces respectively upon cyclic stretch (Figure 1E).

### Rigidity-Dependent Cancer Cells Spread like Normal Cells upon Cyclic Stretch

MDA-MB-231 cells lack the cytoskeletal protein TPM2.1 which is required for proper rigidity sensing (Wolfenson et al., 2016). When TPM2.1 was restored to MDA-MB-231 cells (rigidity-dependent cancer cells) and they were cyclically stretched (5%, 0.5 Hz) for 6 hrs, spread area increased with no change in AR in stark contrast to the wild-type MDA-MB-231 (Figure 2A and S2B) (AR~3 for both non-stretched and stretched MDA-MB-231-TPM2.1 cells). Further, we tested whether cancer cells from another tissue, SKOV3 (human ovarian adenocarcinoma which lacks endogenous TPM 2.1) would behave similarly to MDA-MB-231 cells. SKOV3 cells also demonstrated cyclic stretch-dependent cell elongation (AR~7) after 6 hrs compared to the non-stretched SKOV3 cells (AR~3) (Figure 2B). No increase in the cell area with stretch was observed for SKOV3 cells. However, when TPM2.1 was expressed in SKOV3 cells, they spread more symmetrically (AR~3) and had a 2.5 fold increase in cell area upon stretch when compared with wild-type SKOV3 (Figure 2C). Thus, rigidity-dependent MDA-MB-231 and SKOV3 cells (with restored level of TPM2.1) behaved like wild-type mouse embryonic fibroblasts (MEFs) and breast epithelial cells (MCF10A) upon cyclic stretch (Figure 2D and S4A).

**Figure 2.**
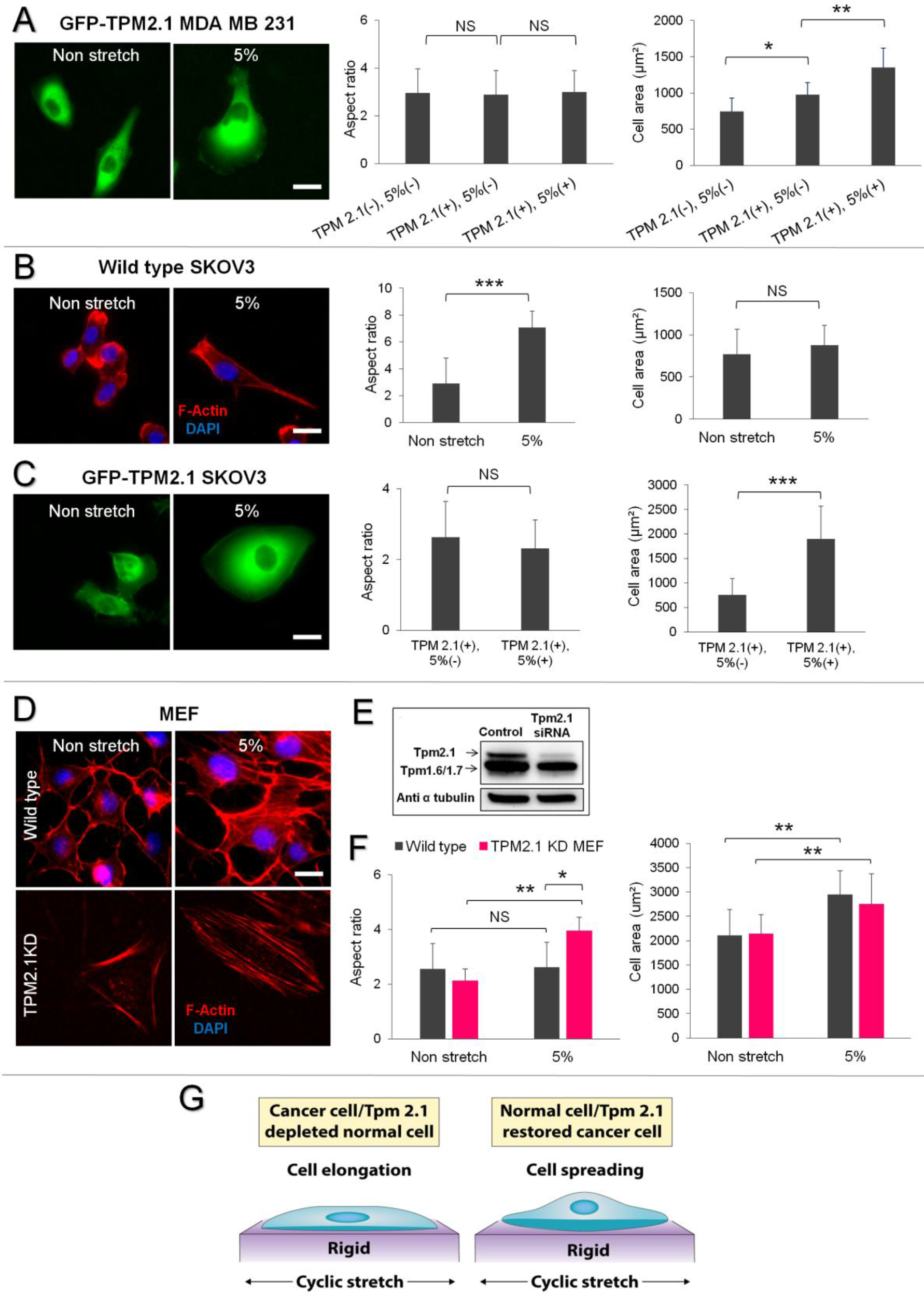
TPM2.1 Restored Cancer Cells Spread upon Cyclic Stretch. **(A)** TPM2.1 restored MDA-MB-231 cells exhibited less elongation (AR~3) and increase in spread area after 6 hrs of cyclic stretch in contrast to wild-type MDA-MB-231, (n=30 cells). **(B)** SKOV3 cells also showed an elongated morphology with 6 hrs of cyclic stretch (AR~7) compared to non-stretched cells (AR~3). No significant increment in the cell area was noticed after cyclic stretching, (n=30 cells). **(C)** TPM2.1 restored SKOV3 cells displayed less elongation (AR~2) and increase in spread area after cyclic stretch, (n=30 cells). **(D)** Representative images showing the morphology of wild-type-and TPM2.1 KD-MEFs on a rigid surface after 6 hrs with and without cyclic stretch. **(E)** Western blots showing the knockdown of TPM2.1 in MEFs. **(F)** Wild-type MEFs showed spread morphology (AR~2) in response to cyclic stretch after 6 hrs, while TPM2.1 KD MEFs were elongated (AR~4). The spread cell area increased for both wild-type-and TPM 2.1KD-MEFs upon cyclic stretch, n=100 cells for aspect ratio and n=30 cells for cell area measurement. (G) Schematic diagram showing the differential effect of cyclic stretch on the morphology of transformed and normal cells. Experiments were repeated at least three times. \**P*< 0.05, \*\**P*< 0.01, \*\*\**P*< 0.001. Scale bar: 20 μm.

To determine if the loss of TPM 2.1, which caused transformation in MEFs and MCF10A, would cause these cells to behave similarly to the transformed cancer cells, we knocked down TPM2.1 in those cells (Figure 2E & S4B). Although wild-type-MEFs and-MCF10A showed a significant increase in spread area with no increase in AR after 6 hrs of cyclic stretching (5%, 0.5 Hz), TPM2.1 KD-MEFs and −MCF10As were elongated upon stretch (Figure 2D and S4A). Statistical analysis confirmed that there was no significant difference in AR values of wild-type cells before and after stretch, whereas TPM 2.1 KD cells demonstrated a two-fold increase in AR after stretch (Figure 2F and S4A). Moreover, there was a stretch-dependent increase in cell area in both wild-type and TPM2.1 KD MEF cells similar to MDA-MB-231 cells after stretch. On the other hand, in MCF10A, we only observed a stretch-dependent cell area increment in wild-type cells. Overall, these results indicate that the cytoskeletal protein TPM2.1 is required for proper sensing of the external mechanical forces that promoted cell spreading in response to the periodic stretching (Figure 2G).

### Cyclic Stretch Reduces Cancer Cell Growth but Stimulates Normal Cell Growth

MDA-MB-231 cells are known for aggressive growth behavior. Therefore, we tested the effect of cyclic stretch on MDA-MB-231, MDA-MB-231-TPM2.1 and normal cell growth (Figure 3A). Unlike previous studies of cyclic stretching effects on fibroblast growth (Cui et al., 2015), cyclic stretching (5%, 0.5 Hz) of MDA-MB-231 cells for 9 hours significantly reduced their proliferation on fibronectin coated rigid and soft surfaces (Figure 3B and 3C). Similarly, cyclic stretch-dependent inhibition of proliferation was observed on the collagen-coated surfaces (Figure S3B). However, a dramatic shift in the effect of stretch on proliferation was found when MDA-MB-231-TPM2.1 cells were stretched. MDA-MB-231-TPM2.1 cells exhibited a higher proliferation rate with cyclic stretch than without on both rigid and soft surfaces. When a similar experiment was repeated using normal cells like MEFs and MCF10A as the control cells, these cells also had a higher proliferation rate (similar to the MDA-MB-231-TPM2.1 cells) with cyclic stretch than without on both surfaces (Figure 3B, 3C and S4C). After MEFs were transformed by TPM2.1 KD, cyclic stretch significantly inhibited proliferation in comparison with the non-stretched cells. Thus, transformation of MEFs by TPM2.1 KD caused a similar behavior to transformed MDA-MB-231 cancer cells with cyclic stretch, indicating that cells in the transformed state had increased sensitivity to stretch-induced growth inhibition. In contrast, normal cells and TPM2.1 expressing MDA-MB-231 cells displayed elevated growth upon cyclic stretch.

**Figure 3.**
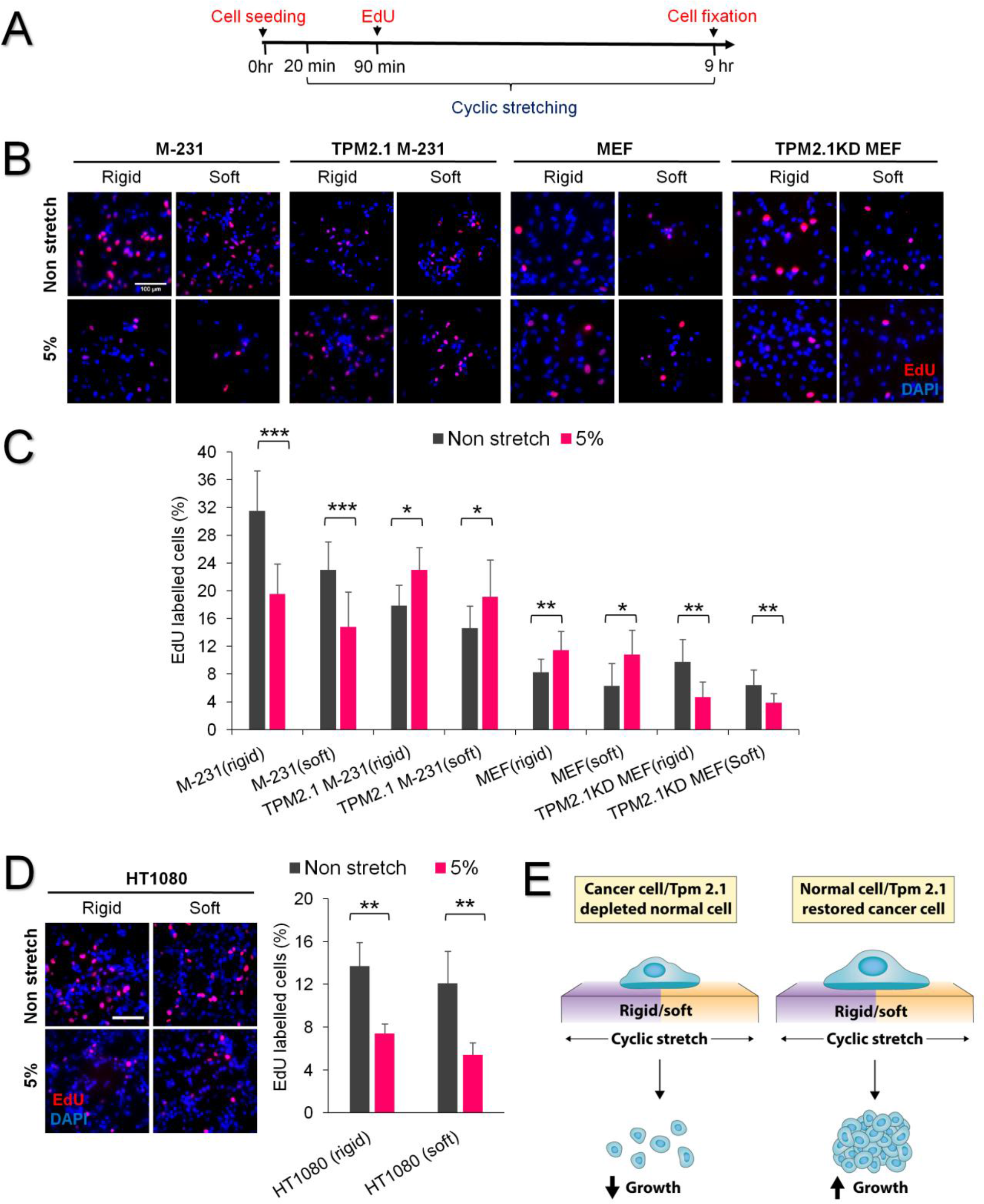
Cyclic Stretch Inhibits Transformed Cell Proliferation. **(A)** Time line depicting the cell proliferation assay protocol. **(B)** Representative images showing the effect of cyclic stretch on the growth of transformed and normal cells (growing cells were labelled by the anti-EdU antibody). **(C)** Cyclic stretch inhibited proliferation of M-231 breast cancer cells on rigid and soft surfaces, while in TPM2.1 restored M-231 cells cyclic stretch increased proliferation. Similarly, cyclic stretching of MEFs (control cells) increased proliferation, whereas cyclic stretch of TPM2.1 KD MEF inhibited cell proliferation, (n>1000 cells for M-231, MEFs, TPM2.1 KD MEFs and n>500 cells for TPM2.1 restored M-231). **(D)** Cyclic stretching of HT1080 fibrosarcoma cells also inhibited proliferation on rigid and soft surfaces, (n>1000 cells). **(E)** Illustration depicting the cyclic stretch-dependent decline in transformed cell growth and increase in normal cell growth. Experiments were repeated at least two times. \**P*< 0.05, \*\**P*< 0.01, \*\*\**P*< 0.001. Scale bar: 100 μm.

To further verify the importance of TPM2.1 in regulating cell proliferation, we analyzed SKOV3 cells (no endogenous TPM2.1). Interestingly, cyclic stretch again caused a reduction in the proliferation of SKOV3 cells on rigid surfaces, while the proliferation rate increased in TPM2.1 expressing SKOV3 cells with cyclic stretch (Figure S5). Thus, transformed cell growth was inhibited by cyclic stretch, whereas in cancer cells that were normalized by TPM2.1 expression, cyclic stretch activated growth.

HT1080 (human fibrosarcoma) cells exhibited normal levels of TPM2.1 but were transformed because of high levels of Tpm3 that competed with TPM2.1 and blocked rigidity sensing (Gateva et al., 2017; Yang et al., 2018). As predicted from other transformed cells results, cyclic stretch decreased the proliferation rate of HT1080 cells on rigid and soft surfaces (Figure 3D). Thus for several different cell lines from different tissues, cyclic stretch inhibited transformed cell growth and promoted normal cell growth (Figure 3E).

### Cyclic Stretch Triggers Cancer Cell Apoptosis but Protects Normal Cells

To determine if inhibition of growth correlated with an increase in cell apoptosis, cells were analyzed for apoptosis after 24 hrs of cyclic stretch by annexin-V immunostaining. As shown in Figure 4A, cyclic stretch activated substantial apoptosis in MDA-MB-231 cells on rigid and particularly on soft surfaces. Closer inspection of apoptotic cells confirmed that they exhibited the characteristics of apoptosis such as membrane blebbing and cell rounding. Interestingly, the apoptosis rate was much higher on the soft surface (34%) where cyclic stretch did not cause cell elongation as compared to the rigid one (18%) (Figure 4B). We observed a similar trend of apoptosis on collagen-coated rigid and soft surfaces (Figure S3C). Further, after transformation of MEFs and MCF10A cells by TPM2.1 KD, cyclic stretching caused a dramatic increase in apoptosis on rigid (20% for MEFs and 54% for MCF10A) and soft (35% for MEFs and 84% for MCF10A) surfaces when compared with the non-stretched control (Figure 4A, 4B and S4D). To strengthen the significance of cyclic stretch in promoting transformed cell apoptosis, MDA-MB-231 cell apoptosis was compared after 24 and 48 hrs of cyclic stretch. We found a notably higher extent of apoptosis (37-39%) after 48 hours on both rigid and soft surfaces (Figure S6) that was likely an underestimate, since many of the cells that died early were washed away. Thus, cyclic stretch caused apoptosis of transformed cells from several different tissues (Figure 4C).

**Figure 4.**
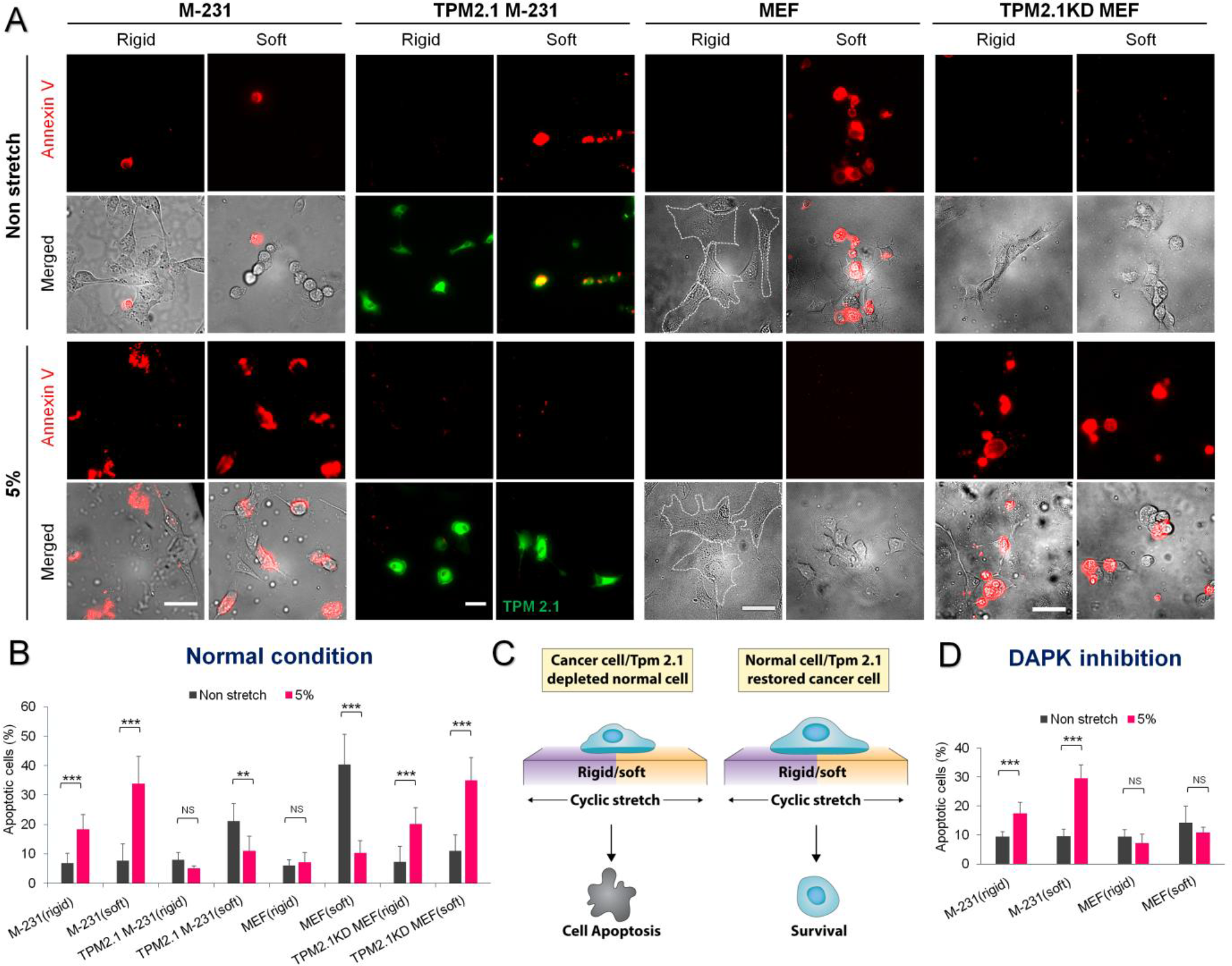
Cyclic Stretch Potentiates Transformed Cell Apoptosis. **(A)** Representative images illustrating the effect of 24 hrs cyclic stretch on apoptosis in M-231, TPM2.1 restored M-231, MEFs and TPM2.1 KD MEFs. Dotted white line indicates location of MEFs. **(B)** Cyclic stretch promoted M-231 cell apoptosis on rigid and soft surfaces. However, TPM2.1 restored M-231 cells experienced negligible apoptosis upon cyclic stretch. Likewise, MEFs (control cells) also demonstrated insignificant apoptosis with cyclic stretch. However, non-stretched MEFs on soft surfaces displayed considerably high apoptosis similar to the non-stretched TPM2.1 restored M-231 cells, indicating cell anoikis on the soft surface. In contrast, TPM2.1 KD MEFs behaved like M-231 cells and experienced an increase in apoptosis after cyclic stretch, (n>350 cells). **(C)** Diagram showing the effect of cyclic stretch on transformed and normal cell apoptosis. **(D)** M-231 cells experienced notably high apoptosis upon stretch even in the presence of DAPK1 inhibitor, suggesting the involvement of DAPK independent apoptotic pathway. However, DAPK1 inhibition rescued the apoptosis in non-stretched MEFs on soft surfaces showing that DAPK1 had a role in normal cell anoikis on soft surface, (n>300 cells). Data are representative of two independent experiments. \**P*< 0.05, \*\**P*< 0.01, \*\*\**P*< 0.001. Scale bar: 50 μm.

To determine if cyclic stretching conditions affected normal cells, the apoptosis assay was performed on rigidity-dependent cells (including TPM2.1 restored cancer cells). When MDA-MB-231-TPM2.1 cells were stretched, there was negligible apoptosis on rigid (6%) or soft (10%) surfaces (Figure 4A and 4B). However, a dramatic increase in apoptosis (21%) was observed in non-stretched MDA-MB-231-TPM2.1 cells on soft surfaces, showing that TPM2.1 restored cancer cells behaved as normal rigidity-dependent cells. To further reconfirm these results, similar experiments were performed on normal cells (MEFs and MCF10As) as the control cells. Both stretched and non-stretched normal cells revealed negligible apoptosis (2-7%) on rigid surfaces (Figure 4A, 4B and S4D). However, on soft surfaces, non-stretched control cells experienced a substantial increase in apoptosis (41-45%), whereas cyclic stretching of the soft surface protected them from apoptosis (5-10%). Thus, cyclic stretching of rigidity-dependent cells inhibited apoptosis, particularly on soft surfaces, whereas cyclic stretching of their transformed counterparts increased apoptosis on soft surfaces.

### Cyclic Stretch Mediated-Cancer Cell Apoptosis Is Not Driven by DAPK1 Activation

Recently, we determined that tumor suppressor Death Associated Protein Kinase1 (DAPK1) causes rigidity-dependent cell apoptosis (anoikis) on soft surfaces but accelerates adhesion assembly on rigid surfaces (Qin et al., 2018). To determine if DAPK1 was involved in stretch-dependent cancer cell apoptosis, we blocked DAPK1 activity with a DAPK1 inhibitor (DAPK1 inhibitor, 100 nM). DAPK1 inhibition decreased MEF apoptosis (14%) on non-stretched soft surfaces (Figure 4D and Figure S7A). As previously reported, we also observed distinct colocalization of DAPK1 with peripheral adhesions not only in stretched and non-stretched normal cells (HFFs) on rigid surface but also in the cells on soft surfaces after cyclic stretch (Figure S7B). In contrast, DAPK1 was cytoplasmic in normal cells on non-stretched soft surfaces. When DAPK1 inhibitor was added to MDA-MB-231 cells during cyclic stretch, there was no decline in apoptosis rate on rigid or soft surfaces indicating that a DAPK1-independent apoptotic pathway was involved in cyclic stretch-induced transformed cancer cell apoptosis.

### Calpain is the Major Downstream Effector of Stretch-Induced Cancer Cell Apoptosis

Since DAPK1 was not involved in cyclic stretch-induced cancer cell apoptosis, we looked for another apoptotic mechanism and found that calpain proteases were implicated in inducing apoptosis in cancer cells (Storr et al., 2011). Calpains were generally activated by a rise in intracellular calcium level and then triggered apoptosis via caspase activity. To test the potential role of calpains in cyclic stretch-induced apoptosis, ubiquitous calpain (calpain 1 & 2) activity was inhibited using the calpain inhibitor (ALLN). Addition of ALLN inhibited apoptosis of MDA-MB-231 cells after cyclic stretch compared to the non-treated counterpart (Figure 5A and S8). Next, to assess the role of specific calpains in cancer cell apoptosis, calpain knock down cells were developed using specific siRNAs. When cyclic stretch experiments were performed on calpain 1 and calpain 2 KD MDA-MB-231 cells, both calpain 1 and 2 KD cells had lower levels of apoptosis than control MDA-MB-231 cells (Figure 5A and S8). However, calpain 2 KD cells showed significantly lower apoptosis, indicating that calpain 2 played the major role in triggering cyclic stretch-induced cancer cell apoptosis (Figure 5B).

**Figure 5.**
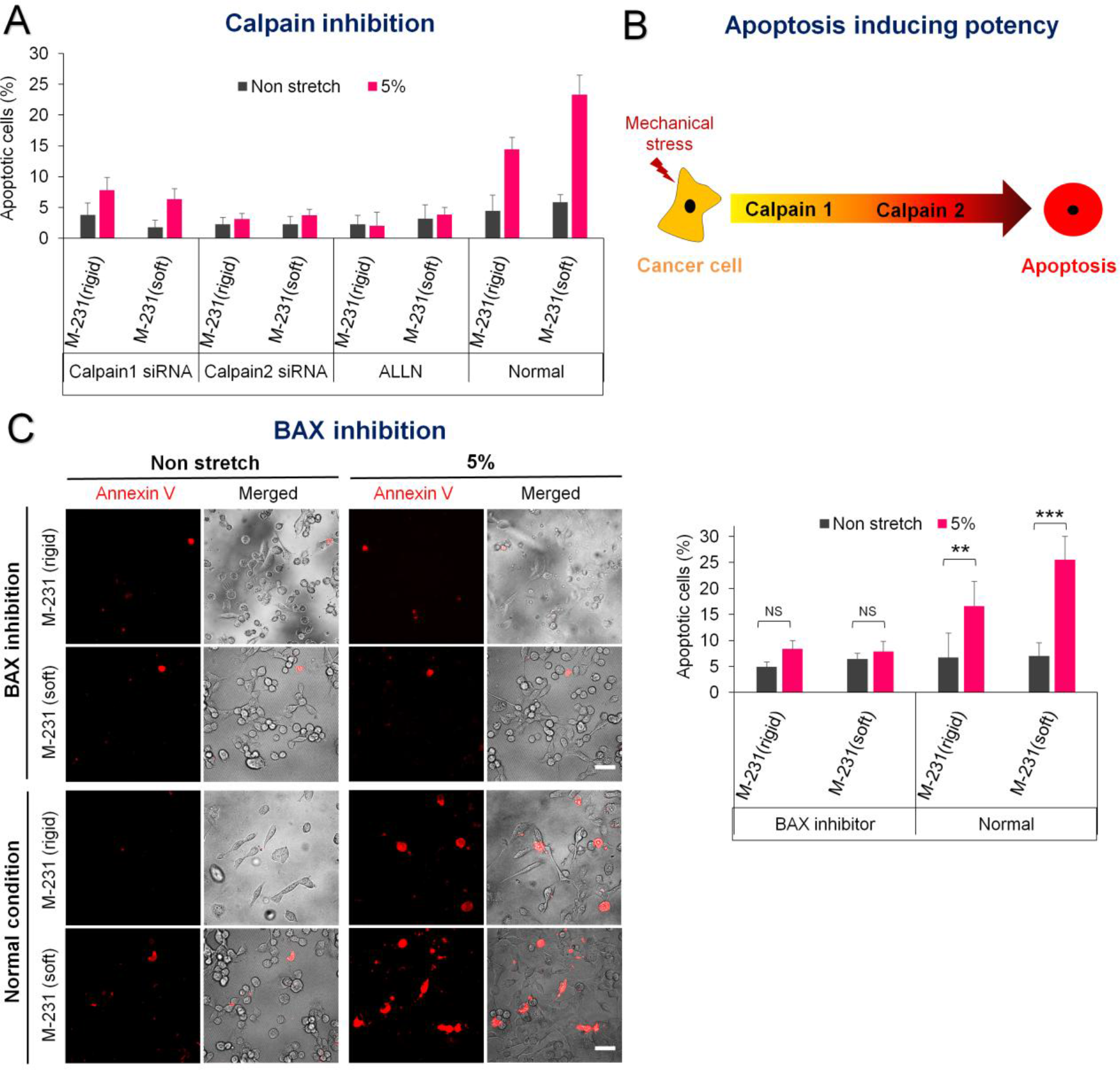
Calpain Acts Downstream of Calcium to Initiate Cyclic Stretch-Induced Apoptosis. **(A)** Cyclic stretch induced M-231 cell apoptosis decreased with Calpain inhibition by using either a specific siRNA against Calpain-1 or −2, or a chemical inhibitor (ALLN). Specifically, Calpain 2 KD caused greater inhibition of cell apoptosis compared to Calpain 1 KD, (n>1000 cells each for Calpain-1 and −2 knockdown assay and n>500 cell for ALLN inhibition assay and control cells). **(B)** Schematic diagram highlighting the major role of Calpain 2 in promoting the cyclic stretch-induced M-231 cell apoptosis. **(C)** M-231 cell apoptosis was further reduced by BAX protein inhibition, implying that BAX protein acts downstream of the calpain protease, (n>300 cells for BAX inhibition assay and n>600 cells for the control). Data are representative of two independent experiments. \*\**P*< 0.01, \*\*\**P*< 0.001. Scale bars: 50 μm.

To understand the downstream effector of calpain cleavage, we focused on the pro-apoptotic molecule BAX which can be cleaved by calpains to induce stress-mediated apoptosis (Wood et al., 1998). Upon activation, BAX molecules translocated from cytoplasm to the mitochondria to initiate the mitochondrial apoptotic pathway (Wolter et al., 1997). To check if BAX was a downstream effector of calpain, MDA-MB-231 cells were stretched in the presence of the BAX inhibitor peptide V5 (blocks mitochondrial translocation of BAX). Interestingly, BAX inhibition reduced cell apoptosis (~10%) in stretched MDA-MB-231 cells (Figure 5C). Thus, it appeared that BAX acted downstream of the calpain protease to initiate the mitochondrial apoptotic pathway.

### Cyclic Stretch-Induced Calcium Influx is the Upstream Effector of Cancer Cell Apoptosis

If stretch-induced apoptosis of transformed cells was caused by calpain, then the cellular level of calcium should increase upon cyclic stretch. Although normal cells have tightly regulated calcium channel expression and regulation (Azimi et al., 2014), greater calcium entry in transformed cells due to cellular stress could have caused cell death via apoptosis (Baig et al., 2016). To determine first if calcium entry was important, we treated cells with the nonspecific calcium channel blocker, Gadolinium, before the application of stretch (Bourne and Trifaro, 1982). Interestingly it blocked the apoptosis in stretched cancer cells (Figure 6A), indicating a possible role of calcium influx as the upstream effector of apoptosis. To actually measure calcium levels during stretch, different cell lines (MDA-MB-231, MDA-MB-231-TPM2.1 and MCF10A) were transfected with a calcium indicator, GECO1 and subjected to the cyclic stretch. In the case of MDA-MB-231 cells, GECO1 intensity increased threefold after 2 hrs of cyclic stretching. (Figure 6B) However, there was no increase in GECO1 intensity in MDA-MB-231-TPM2.1 and MCF10A cells after 2 hrs of cyclic stretching, indicating that stretch triggered-calcium influx did not take place in rigidity dependent-cancer cells and normal cells. Thus, a stretch-induced increase in calcium levels was only found in the transformed cancer cells.

**Figure 6.**
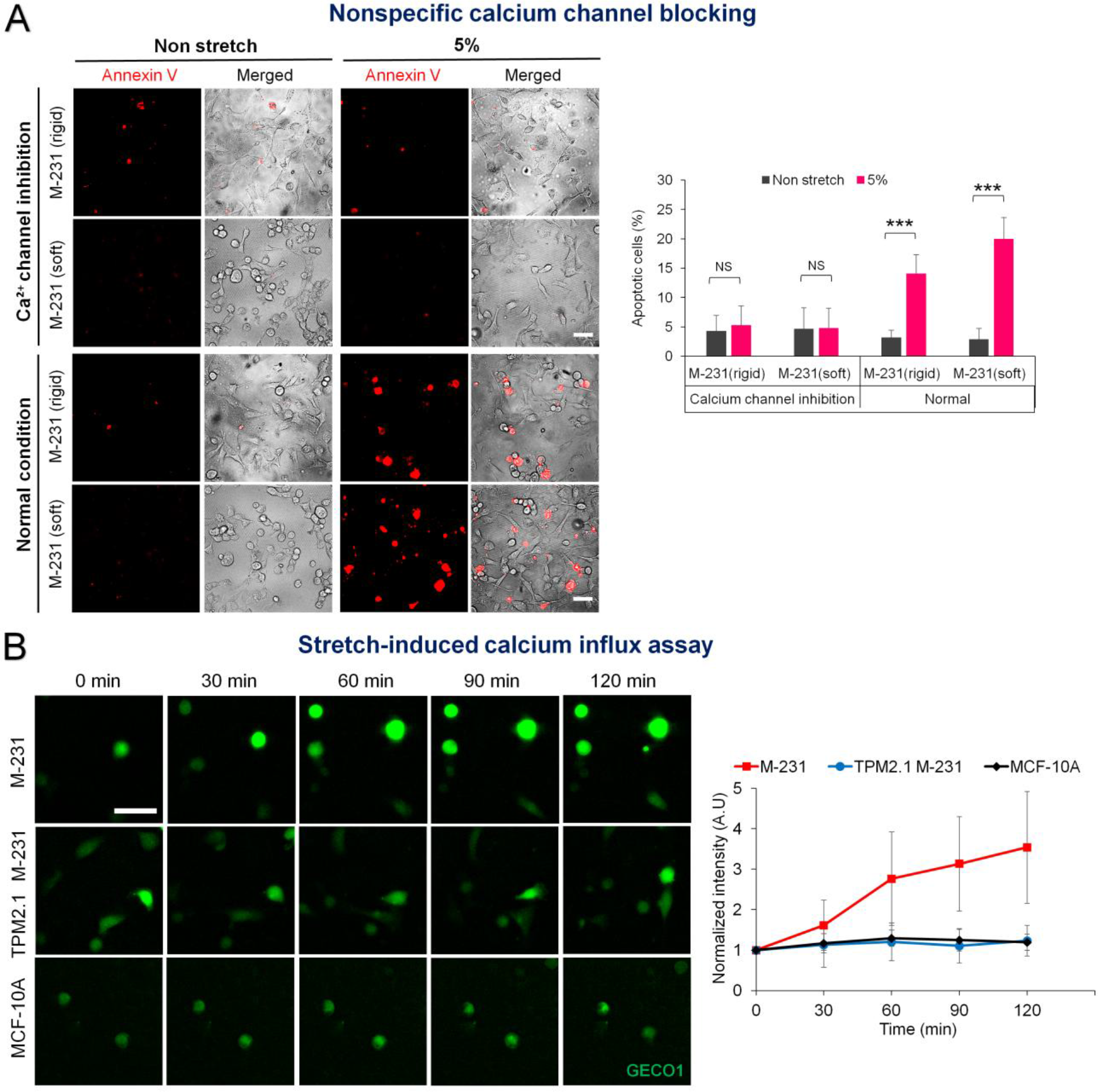
Cyclic Stretch-Mediated Calcium Influx is Upstream Activator of Transformed Cell Apoptosis. **(A)** Cyclic stretch-dependent M-231 cell apoptosis was down regulated by non specific calcium channel blocker, Gadolinium ion, (n>500 cells). Data are representative of two independent experiments. \*\*\*\**P*< 0.001. Scale bar: 50 μm. **(B)** Cells expressing GECO1 calcium indicator were imaged on a cyclically stretched rigid surface. Representative time-lapse montages displaying the progressive increase in GECO1 intensity in M-231 cells. In stark contrast, no such increase in GECO1 intensity was observed in TPM2.1 restored M-231 cells and normal MCF10A cells. After 2 hrs, the GECO1 intensity increased by three-fold in M-231 cells compared to other groups, n>15 cells from three independent experiments were used. Scale bar: 50 μm.

### Mechanosensitive Piezo1 Channels Promote Cyclic Stretch-Induced Cell Apoptosis

The linkage between stretch-induced apoptosis and calcium entry implied the involvement of mechanosensitive calcium channels. Piezo channels were activated by mechanical perturbations of cells and enabled calcium entry to trigger intracellular calcium dependent signaling in many physiological processes (Coste et al., 2010). We therefore compared the level of apoptosis in Piezo1 expressing HEK 293 cells (transformed human embryonic kidney cells) and Piezo1 knock out HEK 293 cells (piezo KO HEK 293) after cyclic stretch on rigid surfaces for 6 hrs. Cyclic stretch caused an increase in the apoptosis of piezo expressing HEK 293 cells, while negligible apoptosis was observed in piezo KO HEK 293 cells (Figure 7A). Transformed HEK 293 cells lack endogenous TPM2.1. When the apoptosis assay was performed on TPM2.1 restored HEK 293 cells either with or without Piezo1, negligible apoptosis was noticed in piezo KO HEK 293 or Piezo1 expressing HEK 293 cells with TPM2.1 restored (Figure 7B and Figure S9). Thus, we suggest that stretch-induced transformed cell apoptosis involved Piezo1 channels as well as the transformed state.

**Figure 7.**
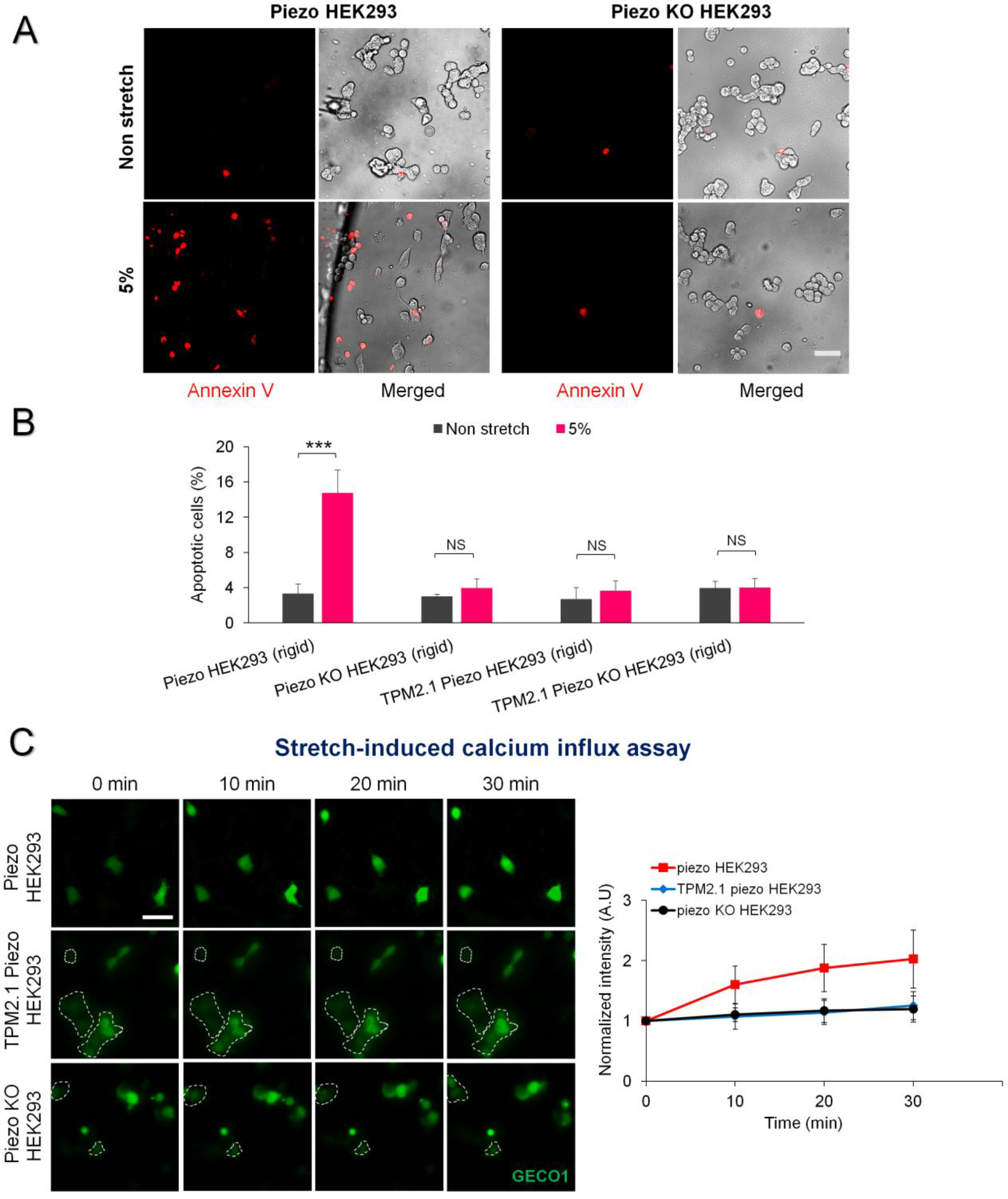
Mechanosensitive Piezo1 Channels Needed for Cyclic Stretch-Induced Apoptosis. **(A)** Representative images illustrating elevated level of apoptosis in piezo expressing transformed HEK293 cells upon cyclic stretch compared to piezo KO HEK 293 cells. **(B)** Piezo expressing HEK293 cells exhibited significant apoptosis (~15%) upon cyclic stretch compared to piezo KO HEK 293 and TPM2.1 restored HEK 293 either with or without Piezo1 (~4% in all cases), (n>300 cells). Data are representative of two independent experiments. \*\*\**P*< 0.001. Scale bar: 50 μm. **(C)** Cells expressing GECO1 calcium indicator were monitored on a cyclically stretched rigid surface. Representative time-lapse montages show a progressive increase in GECO1 intensity in Piezo expressing HEK293 cells. No such increase in GECO1 intensity was observed in TPM2.1 restored HEK293 and Piezo KO HEK293 cells. Two-fold increase in the GECO1 intensity was observed in Piezo expressing HEK293 cells after 30 min compared to other groups, n=16 cells from three independent experiments were used. Scale bar: 50 μm.

To determine if the rise in cellular calcium level relied upon Piezo1 channels, cyclic stretch was applied on GECO1 transfected Piezo1-and Piezo1 KO-HEK 293 cells and TPM2.1 restored Piezo1 HEK 293 cells. Of these three different cells, only piezo1 expressing HEK 293 cells showed a twofold increase in GECO1 intensity within 30 min on rigid surface, while rigidity-dependent or Piezo1 KO cells did not display any significant increase in the intensity (Figure 7C). All together, these findings confirmed that stretch-induced calcium entry and subsequent apoptosis in transformed HEK 293 cells depended upon Piezo1 channels.

## Discussion

These results indicate that transformed cells exhibit cyclic stretch-induced apoptosis regardless of the tissue origin, whereas rigidity-dependent normal cells show cyclic stretch-induced growth and survival. For several different cell lines, the presence of TPM2.1 causes rigidity-dependent growth, while depletion of TPM2.1 results in transformed growth despite different tissue backgrounds. In transformed cells, cyclic stretch initially causes cell elongation on rigid surfaces that is dependent on the stretch frequency as well as the magnitude of strain. The frequency and magnitude of stretch were in a physiologically relevant range for exercise and those parameters were used in all further studies. Longer period of cyclic stretch inhibits transformed cell growth and increases apoptosis, particularly on soft surfaces. In contrast, cyclic stretch promotes normal cell growth and inhibits apoptosis, particularly on soft surfaces. We find that this relationship holds for cells from breast, ovaries, kidneys, connective tissue and skin. Further, the stretch-induced apoptosis is calpain dependent (primarily calpain 2) and calpain acts downstream of stretch-mediated calcium influx. The mechanosensitive Piezo1 calcium channels are needed for calcium influx in the background of the transformed cell state.

Numerous studies regarding the transformed cell state imply that normal cells from a variety of different tissues can be transformed. During the cell transformation, there is a shift from an epithelial to mesenchymal phenotype in many cases; but the transformed phenotype is strongly correlated with the loss of rigidity sensing and a change in the cortical actin organization that makes cancer cell softer (Lin et al., 2015; Wolfenson et al., 2016). However, the traction forces that most cancer cells exert on matrices are higher than their normal counterparts and restoration of rigidity sensing in those cancer cells causes a decrease in traction forces (Northcott et al., 2018; Wolfenson et al., 2016; Yang et al., 2018). Thus, it is logical to consider the ‘transformed state’ as a distinct cell state that can occur in cells from widely different tissues with different expression patterns. Further, cells from different tissues can be toggled between the transformed and the normal state by changing the level of expression of several cytoskeletal proteins that enable or block the matrix rigidity sensing (Yang et al., 2018).

A relatively common feature of transformed cancer cells is that they have elevated levels of calcium channels and calpain protease (Azimi et al., 2014; Storr et al., 2011). The higher level of a mechanosensitive calcium channels and their altered functioning could explain the increased susceptibility of transformed cancer cells to stretch-induced apoptosis. However, the transformation of fibroblasts and epithelial cells with TPM2.1 KD will also sensitize those cells to stretch-induced apoptosis. This indicates that transformed cells in general, whether from a tumor or from a normal cell background, will undergo apoptosis in response to the appropriate mechanical forces that can be mimicked by cyclic stretch.

There are previous reports of mechanical force-induced cancer cell death. Continuous flow forces can cause the apoptosis of several different cancer cells attached to surfaces, whereas oscillatory flow forces do not (Lien et al., 2013). The apoptotic pathway involves bone morphogenetic protein receptor, Smad1/5, and p38 MAPK unlike our findings. Another study shows that high shear forces on suspended circulating tumor cells initiates apoptosis possibly through an oxidative stress-induced mitochondrial apoptotic pathway (Regmi et al., 2017). Further, recent findings reveal that the stretching of mice for ten minutes a day for four weeks suppresses the breast cancer growth by 50% compared to the non-stretched controls (Berrueta et al., 2018). From these and other studies, there are strong indications that transformed cancer cells may be mechanically vulnerable and thus sensitive to mechanical forces. Here we show that the increased mechanical sensitivity is linked to the transformed cell state and not to a specific organ or a specific cell type. Thus, it seems that this feature could be exploited to damage many different types of tumor cells, particularly in a metastatic state where cells are not protected by tumor fibrosis.

In Figure 8, we present a working model of mechanical stretch-induced transformed cancer cell death, which we are describing as ‘mechanoptosis’. Prolonged application of mechanical stresses on the transformed cells triggers a rise in calcium levels through altered calcium homeostatic mechanisms, most likely increased Piezo1 activity. The rise in intracellular calcium level causes activation of calpain 2 protease which initiates a mitochondrial apoptotic pathway through its downstream effector, the BAX molecule. In contrast, mechanical stretching of normal cells does not cause a rise in calcium levels because of a more tightly regulated calcium homeostatic mechanism. Thus, cyclic stretch promotes the normal cell growth and survival especially on the soft surfaces.

**Figure 8.**
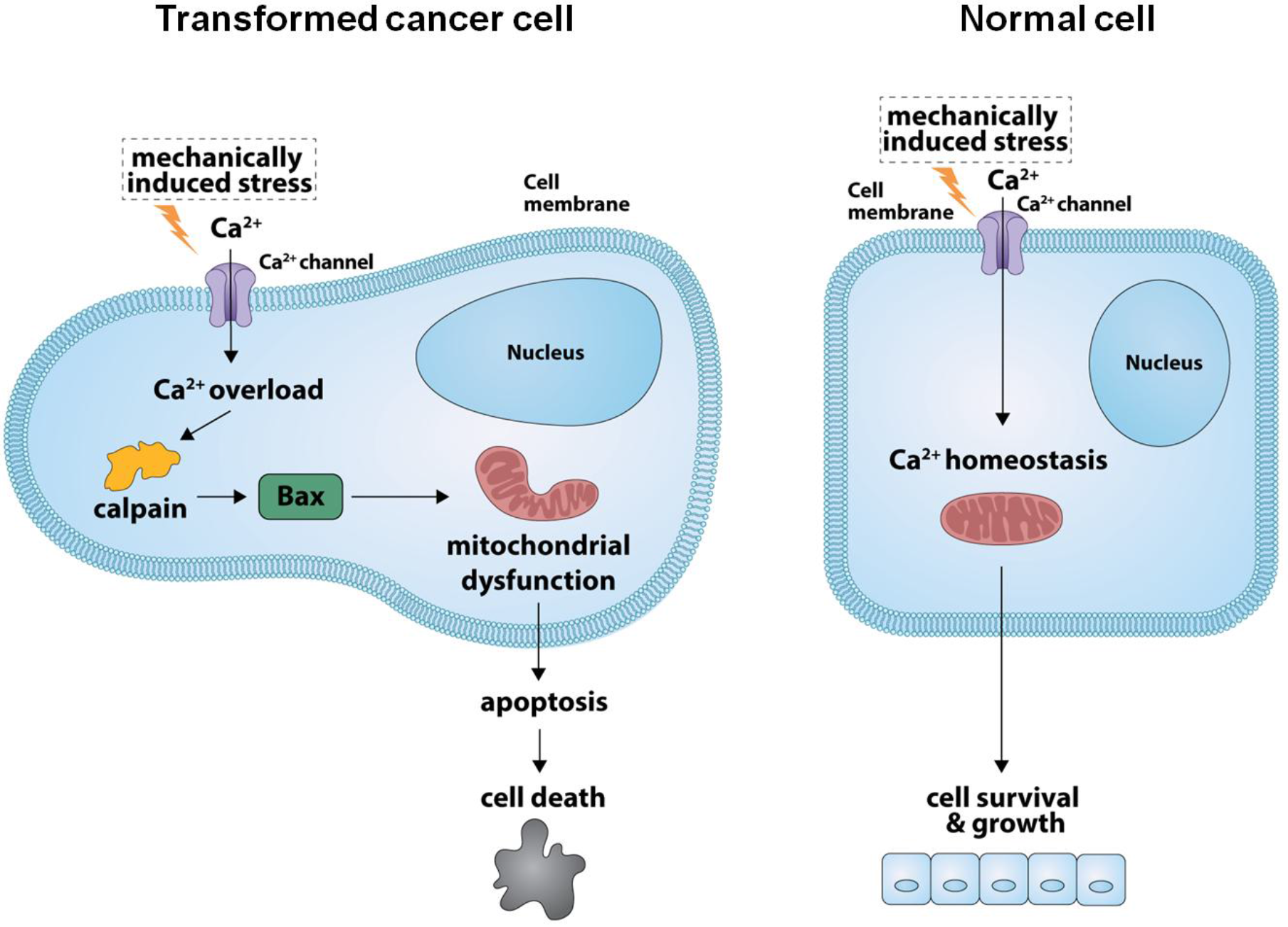
Proposed Model of Transformed Cell Mechanoptosis. Prolonged cyclic stretching of transformed cells leads to intracellular calcium overloading, activation of calpain protease and in turn initiation of the mitochondrial apoptotic pathway through calpain downstream effector, BAX molecule. In contrast, mechanical stretching of normal cells does not trigger calcium influx through their tightly regulated calcium channels. In fact, it promotes the normal cell growth and survival through normal calcium homeostasis.

Although most cancer cells are transformed, there are many other aspects that are important for tumor growth (Hanahan and Weinberg, 2000; Hirashima et al., 2013; Nishida et al., 2006). Nevertheless, the transformed cell state appears to be associated with changes in important cell functions such as the loss of rigidity sensing. Many other factors are also altered beyond the loss of rigidity sensing in transformed cells, making it difficult to link the increased mechanical sensitivity to a single factor. Despite the fact that we do not fully understand why transformed cells are more sensitive to calcium loading with cyclic stretch, that sensitivity can provide new approaches for killing cancer cells while actually stimulating the normal cell growth. Hence, much more work is needed to determine if this feature can be exploited in an in vivo setting to damage transformed cancer cells.

## Material and Methods

### Cell Culture

All cell lines MDA-MB-231(gift from Dr. Jay Groves, MBI, NUS), SKOV3 (gift from Dr. Ruby Huang, CSI, NUS), HT1080 (ATCC), human foreskin fibroblasts (HFF, ATCC), transformed human embryonic kidney cells (piezo expressing HEK 293 and piezo knock-out HEK 293, gift from Boris Martinac, VCCRI Sydney and Ardem Patapoutian) and mouse embryonic fibroblasts (MEF, ATCC) were cultured in high glucose DMEM (Thermo Fisher Scientific) supplemented with 10% FBS and 1% penicillin and streptomycin at 37°C in 5% CO2 environment. MCF10A, human breast epithelial cells (ATCC) were cultured in specialized growth medium as mentioned before (Wolfenson et al., 2016). For cell experiments, high-glucose DMEM containing 10% FBS and 1% penicillin and streptomycin was used. Cells were trypsinized using TrypLE (Thermo Fisher Scientific) and cultured on human plasma fibronectin (10 μg/ml, Roche) or rat tail collagen-I (20 μg/ml, Sigma) coated PDMS surface. Pharmacological inhibitors used are as follows: DAPK1 inhibitor (100 nM, EMD Millipore), calpain inhibitor ALLN (100 μM, Sigma Aldrich), non-specific calcium channel blocker gadolinium chloride (20 μM, Sigma Aldrich) BAX inhibitor peptide V5 (200 μM, Merck). For inhibition assays, inhibitors were added to the medium one hour post cell seeding and cyclic stretching was applied one to two hours after addition of inhibitors. In general, cyclic stretching was applied 20 min after the cell seeding on the cell stretching post (unless otherwise stated).

### Plasmids and Transfection

For cell transfection using GFP tagged-TPM2.1 plasmid and RFP tagged-GECO1 calcium indicator, Neon electroporation system (Invitrogen) was used according to the manufacturer’s protocol. To perform TPM2.1 knockdown, cells were plated in a six well plate. TPM2.1 siRNA (Qiagen) transfection was performed on the next day using lipofectamine RNAiMAX (Invitrogen) according to the manufacturer’s protocol. Calpain siRNAs were prepared by Protein Cloning Expression Facility, MBI, NUS using following siRNA sequences adapted from previous reports. The calpain siRNA primers were: Calpain 1 primer (5’-AAGAC CUAUGGCAUCAAGUGG-3’; 5’-AAGAAGUUGGUGAAGGGCCAU-3; ‘5’-AAGCUAGU GUUCGUGCACUCU-3’; 5’-AAG AGGAGAUUGACGAGAACU-3’ (Upla et al., 2008; Wu et al., 2006) and Calpain 2 primer (5’-AAGACUUCACCGGAGGCAUUG-3’; 5’-AAGAUG GGCGAGGACAUGCAC-3’) (Ma et al., 2012; Xu et al., 2010). Calpain knockdown cells were generated by cell transfection using lipofectamine RNAiMAX.

### Pillar and Flat Surface Fabrication

Pillar substrates were prepared with PDMS (10:1 prepolymer to curing agent ratio, Sylgard 184; Dow Corning) spin coating on plastic molds at 1800 rpm for 1 min and cured at 80°C for 3 hrs. The PDMS membrane was then peeled off the mold to serve as the surface for cell culture as shown in Figure S1A. Pillars were fashioned in a square grid with 0.5 μm diameter, 1.8 μm (~8kPa, soft) or 0.8 μm (~55kPa, rigid) height and 1.5 μm centre-to-centre distance among pillars (Lohner et al., 2018). The flat PDMS layer (2 MPa rigid, 60-70 μm thick) was prepared by spin coating PDMS (10:1 prepolymer to curing agent ratio) on a plastic sheet at 1800 rpm for 1 min, cured at 80°C for 3 hrs and peeled off to treat as the loading surface for cell culture.

### Stretching Device Fabrication and Operation

The scheme of stretching device fabrication was shown in Figure S1A. Standard lithography technique was used to fabricate the PDMS stretching device. Briefly, PDMS solution (10:1 prepolymer to curing agent ratio) was spin coated on a silicon wafer containing microfeatures at 500 rpm for 30 sec and cured at 80°C for 3 hrs. PDMS layer embossed with microfeatures was peeled off, fixed to the glass bottom dish and subsequently covered with another thin PDMS layer (60-70 μm thick). Glycerol (90%) was used as a lubricant to avoid sticking of thin PDMS layer to the lower cell loading post. A PDMS cube with hole in middle was mounted on the top of thin PDMS layer as shown in Figure S1A. Finally, the PDMS cube was connected to the pump assembly. As depicted in Figure S1B & S1C, negative pressure was generated in the hollow ring surrounding cell loading post by a pump which pulled the thin PDMS membrane down to create membrane stretching. Different negative pressures were applied to measure the magnitude of the strain generated at each pressure and the percent strain was then characterized by measuring the changes in centre to centre distance between the pillars (Figure S1D & S1E). To control the frequency of cyclic stretch, the Labview program was used.

### Proliferation Assay

Cells were serum starved overnight to synchronize the cell cycle prior to cell seeding. EdU (5-ethynyl-2’-deoxyuridine) reagent kit (Invitrogen) was used according to the manufacturer’s instruction. The proliferation assay was performed after 9 hrs.

### Apoptosis Assay

To assess cell apoptosis, Annexin V-Alexa Fluor 488 or Annexin V-Alexa Fluor 594 conjugates (Thermofisher Scientific) were used according to the manufacturer’s protocol. Cells were incubated with Annexin V solution after the specified time mentioned in the results section.

### Immunocytochemistry and Fluorescence Microscopy

Cell samples were fixed with 4% paraformaldehyde (Thermofisher Scientific) in PBS for 10 min and then permeabilized with 0.5% Triton X-100 for 5 min. Bovine Serum Albumin solution (2%) (BSA) was used as a blocking buffer and samples were treated with BSA for 1 hr. Samples were then incubated with primary antibodies of rabbit polyclonal anti DAPK1 (Abcam, 1:200), mouse monoclonal anti Paxillin (BD Bioscience, 1:400) and mouse monoclonal anti tropomyosin (1:200, Merck) for 1 hr at 37°C followed by treatment with Alexa Fluor-488 and −594 secondary antibodies (Invitrogen). For F-actin staining, cells were incubated with Alexa Fluor 594 phalloidin antibody (Invitrogen) for 1 hr. DAPI dye (Merck) was used to stain cell nucleus. Fluorescence and bright field images were acquired using Delta Vision System (Applied Precision) on an Olympus IX70 microscope and equipped with CoolSNAP HQ2 CCD camera (Photometrics, Tucson, USA). Live cell imaging was done using same Delta Vision System maintained at 37°C, 5% CO2 condition. Optical images were acquired with an EVOS digital fluorescence microscope (Fisher Scientific).

### Western Blot

The cells were harvested after at least 48 hrs with the respective siRNA’s, pelleted, washed with PBS and then lysed with RIPA buffer (Sigma-Aldrich) supplemented with the 1X cOmplete Protease Inhibitor cocktail tablet (Roche, Cat no: 4693116001). The lysed cell mixture was centrifuged at 15000 rpm, 4°C for 30 mins. The supernatant was collected and mixed with 2X loading dye (2X Lammeli Buffer, Bio-Rad, Cat no: 1610737) + Beta-Mercaptoethanol (Sigma-Aldrich). The samples were then denatured for 10 mins at 95°C and run on a 4-20% Mini-PROTEAN ^®^ TGX precast protein gels (Bio-Rad, Cat no: 4561094). The gels were then transferred onto membranes. The membranes were blocked with 5% BSA solution in 1X TBST (Tris-Buffered Saline with Tween-20) for 1 hr and incubated with primary antibody overnight at 4°C. The membranes were washed three times in TBST (10 mins per wash) followed by secondary antibodies in 1XTBST (Horse Radish Peroxidase -HRP) treatment for 1 hr. The membranes were again washed three times in 1X TBST. The chemiluminescence of the membranes was developed using Super Signal Femto Substrate Kit (Pierce) and developed using ChemiDoc Touch Imager (Bio-Rad). Primary antibodies and conditions used are as follows: Calpain 1 antibody (Rabbit, 1:1000, Abcam ab39170), Calpain 2 antibody (Rabbit, 1:1000, Abcam ab39165), TPM 2.1 (Mouse, 1:1000, Merck), GAPDH (Mouse, 1:3000, Abcam ab8245). The secondary antibodies, HRP-conjugated goat anti-rabbit IgG (bio-Rad 170-6515) and goat anti-mouse IgG (bio-Rad 170-6516) were used at half of the primary antibody concentration.

## STATISTICAL ANALYSIS

Statistical difference was calculated using two-tailed Student *t*-test. *P* value <0.05 was considered statistically significant. All data represented as mean ± standard deviation.

## Supporting information

## ACKNOWLEDGEMENTS

We would like to thank all group members of Sheetz lab. A.T and M. Y are supported by the Singapore Ministry of Education Academic Research Fund Tier 3 (MOE grant No: 2016 T3-1-002). M.S is supported by NIH grants, NUS grants and Mechanobiology Institute, National University of Singapore.

## AUTHOR CONTRIBUTION

M.S and A.T designed study; A.T performed cell experiments, device fabrication and data analysis; A.T and M.Y performed calcium indicator experiments; Y.W fabricated molds; Y.N and C.T.L engineered stretching device; A.H developed western blots; A.T and M.S wrote manuscript. All authors read manuscript and commented on it.

