## Supplementary material for "Mechanical Stretch Kills Transformed Cancer Cells"

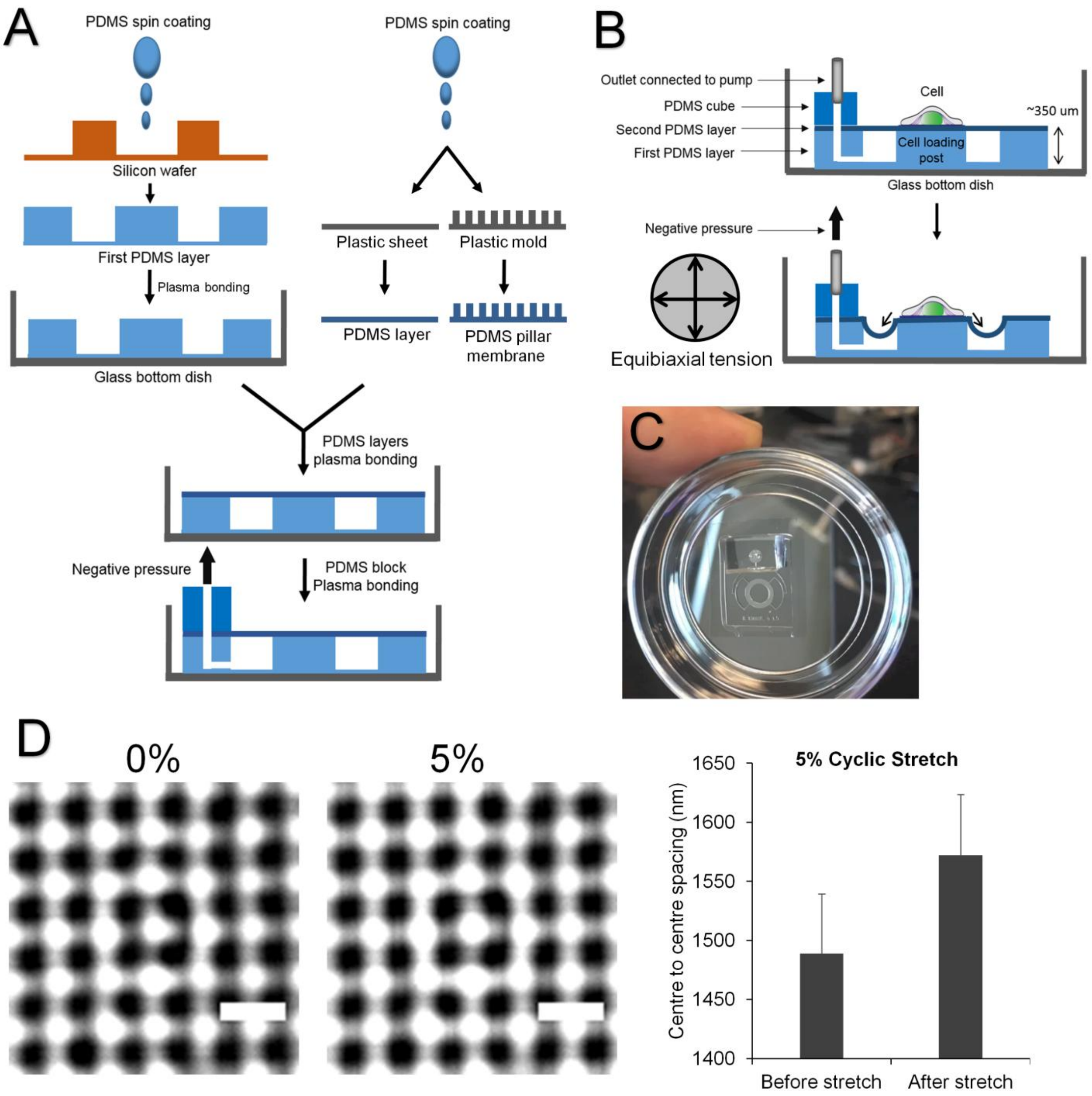


**Figure S1. Stretching Device Fabrication, Working and Characterization**

**(A)** The scheme of stretching device fabrication. Standard lithography technique was used to fabricate the PDMS stretching device. **(B)** The schematic depicting the working of stretching device. **(C)** Picture of the actual stretching device mounted on glass bottom dish. **(D)** Stretching device was covered with soft pillar membrane. Percent strain was calculated by measuring the change in centre to centre distance between pillars, n=200 pillars, scale bar: 2 µm.

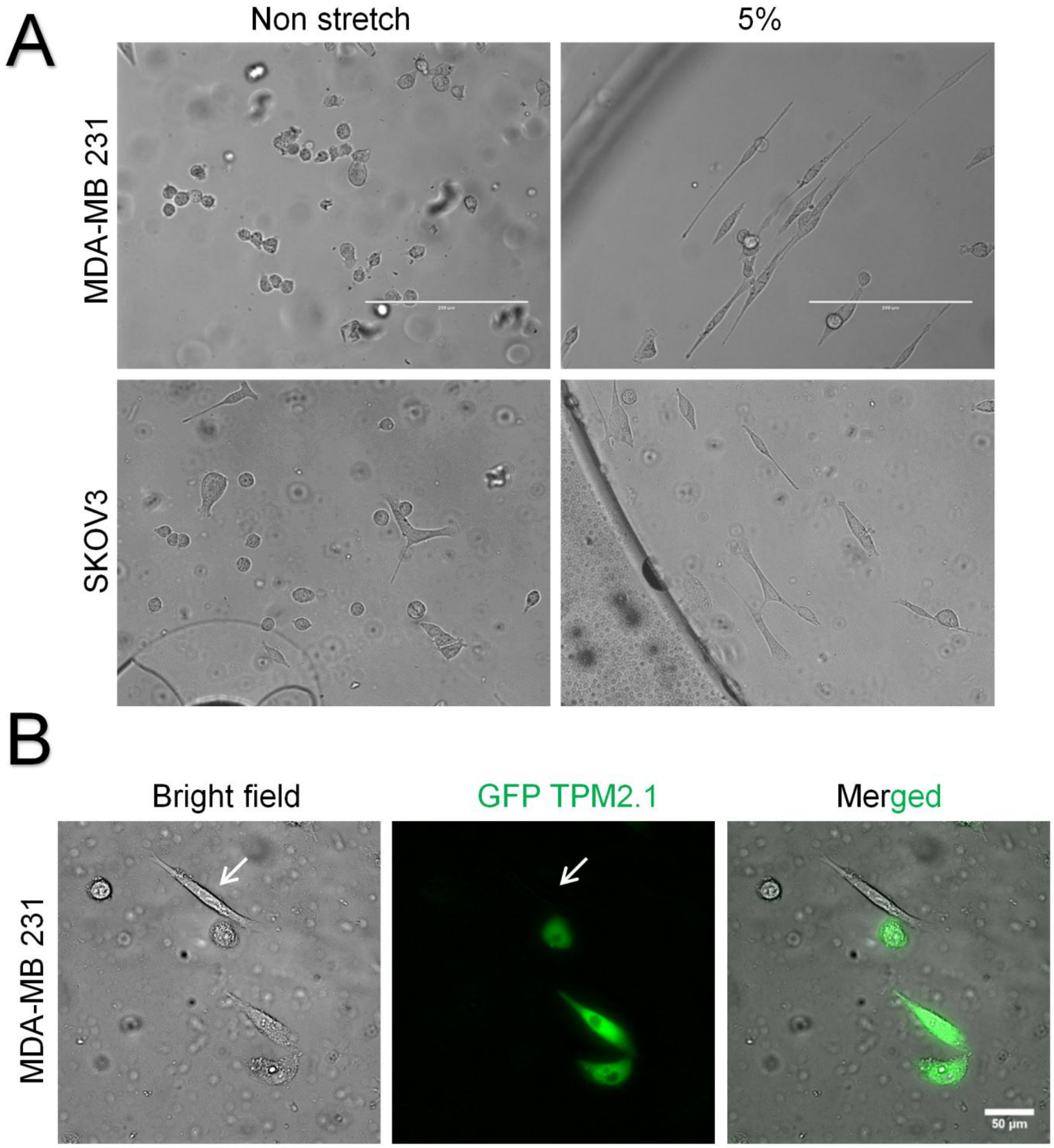


**Figure S2. Cancer Cells Elongate upon Cyclic Stretch, while TPM2.1 Restored Cancer Cells Spread**

**(A)** Representative images of M-231 and SKOV3 cancer cells displaying elongated morphology after 6 hrs of cyclic stretch (5%, 0.5Hz), scale bar: 200 µm. **(B)** TPM2.1 transfected M-231 cells exhibited spread morphology compared to wild-type M-231 cells. White arrow indicates non-transfected cancer cell showing an elongated morphology, scale bar: 50µm.

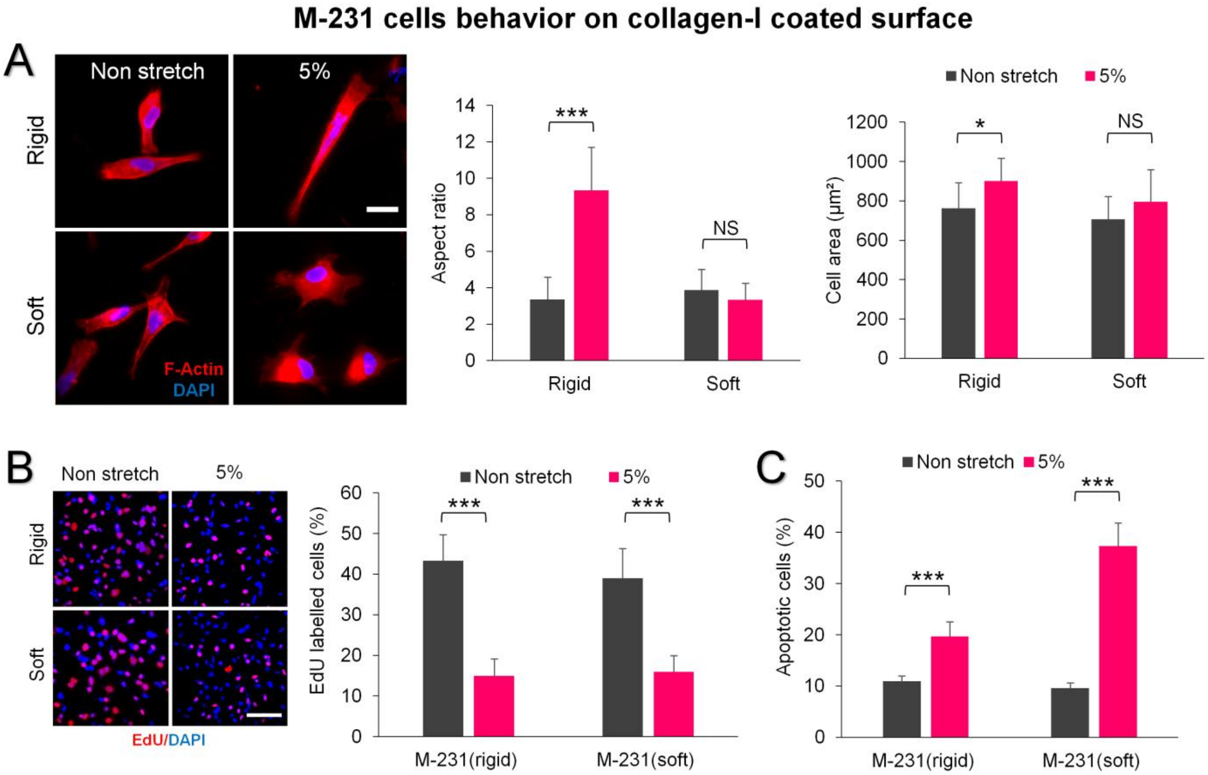


**Figure S3. Effect of Collagen-I Coated Surface on M-231 Cancer Cell Morphology, Proliferation and Apoptosis after Cyclic Stretching**

**(A)** M-231 cancer cells showed elongated morphology (AR~9) on collagen-I coated rigid surfaces upon cyclic stretch (5%, 0.5 Hz) after 6 hrs, while such cell elongation was absent on the soft surface. Stretch dependent increase in the cell area was observed on rigid surface only, n=100 cells for cell aspect ratio and n=30 cells for cell area measurement. Experiments were repeated at least two times, scale bar: 20 μm. **(B)** Cyclic stretching of cells for 9 hrs caused significant reduction in M-231 cell proliferation on collagen coated rigid and soft surfaces, n>1000 cells. Experiments were repeated at least two times, scale bar: 100 μm. **(C)** Apoptosis rate was also found notably high on collagen coated rigid (20%) and soft (37%) surfaces after 24 hrs of cyclic stretch, n>250 cells. Experiments were repeated at least two times. *$P < 0.05$, **$P < 0.01$, ***$P < 0.001$.

### MCF10A cells study

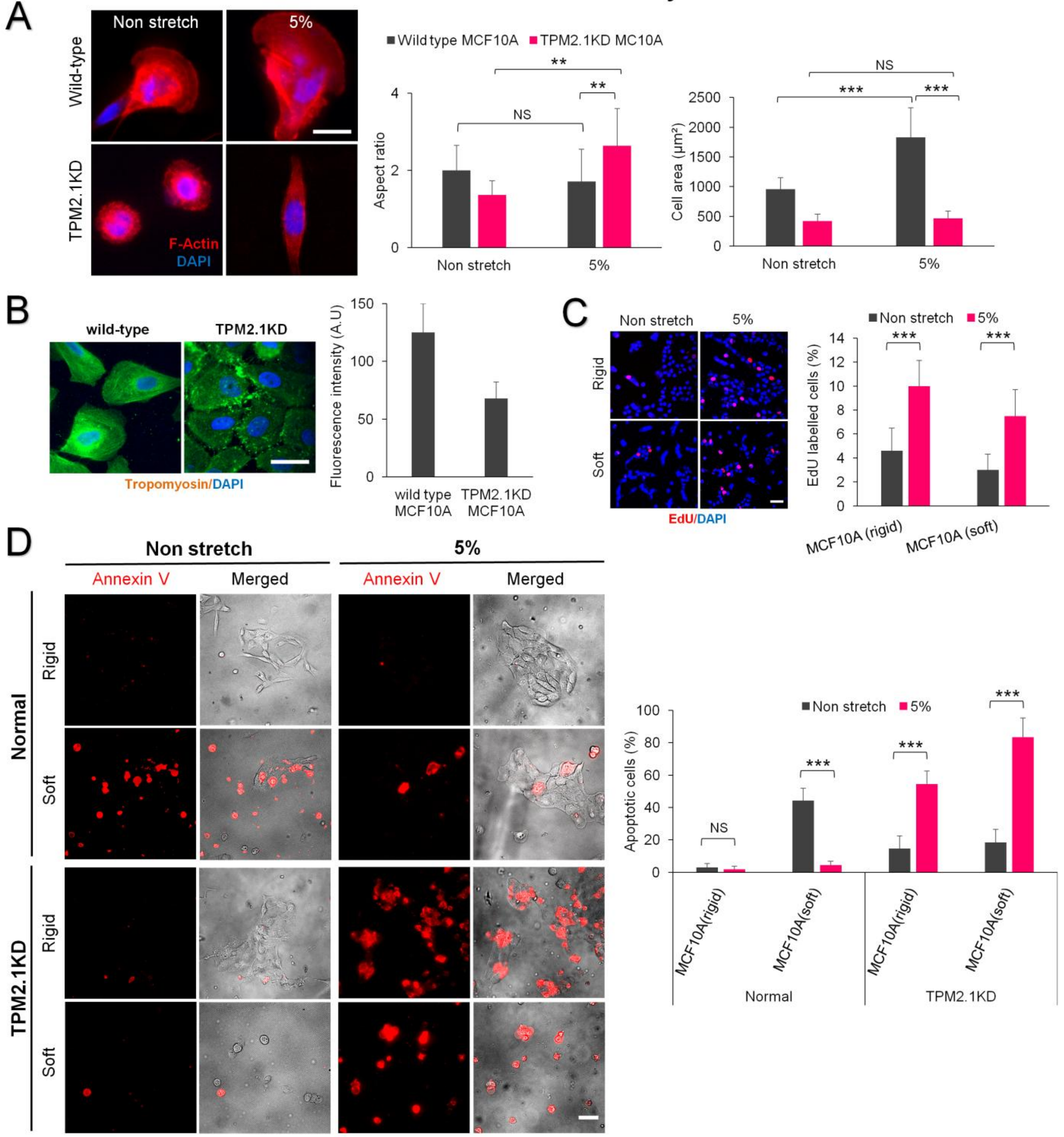

**Figure S4. Effect of Cyclic Stretch on MCF10A Cell Morphology, Proliferation and Apoptosis (A)** Representative images showing the morphology of wild-type- and TPM2.1 KD-MCF10A cells on rigid surface after 6 hrs with and without stretch. Wild-type MCF10A cells showed spread morphology (AR~2) upon stretch, while TPM2.1 KD MCF10A cells displayed elongated morphology (AR~2.6) compared to non stretched counterpart (AR~1.3). Cell area increment upon stretch was observed only for wild-type MCF10A, n=30 cells for aspect ratio and cell area quantification. Experiments were repeated at least two times, scale bar: 20 µm. **(B)** Representative images showing tropomyosin expression in wild-type- and TPM2.1 KD-MCF10A cells. Fluorescence intensity measurement confirmed substantial decrease in tropomyosin expression in TPM2.1 KD cells, (n>25 cells), scale bar: 20 µm. **(C)** Cyclic stretch promoted MCF10A cell proliferation on rigid and soft surfaces, n>900 cells. Experiments were repeated at least two times, scale bar: 50 µm. **(D)** Representative images illustrating the impact of 24 hrs cyclic stretching on wild-type MCF10A and TPM2.1 KD MCF10A cell apoptosis. Wild-type MCF10A cells demonstrated negligible apoptosis with cyclic stretch on both surfaces. On contrary, notably high apoptosis was found in TPM2.1 KD MCF10A cells upon cyclic stretch, (n>500 wild-type MCF10A cells and n>250 TPM2.1 KD MCF10A cells), scale bar: 50 µm. *$P< 0.05$, **$P< 0.01$, ***$P< 0.001$.

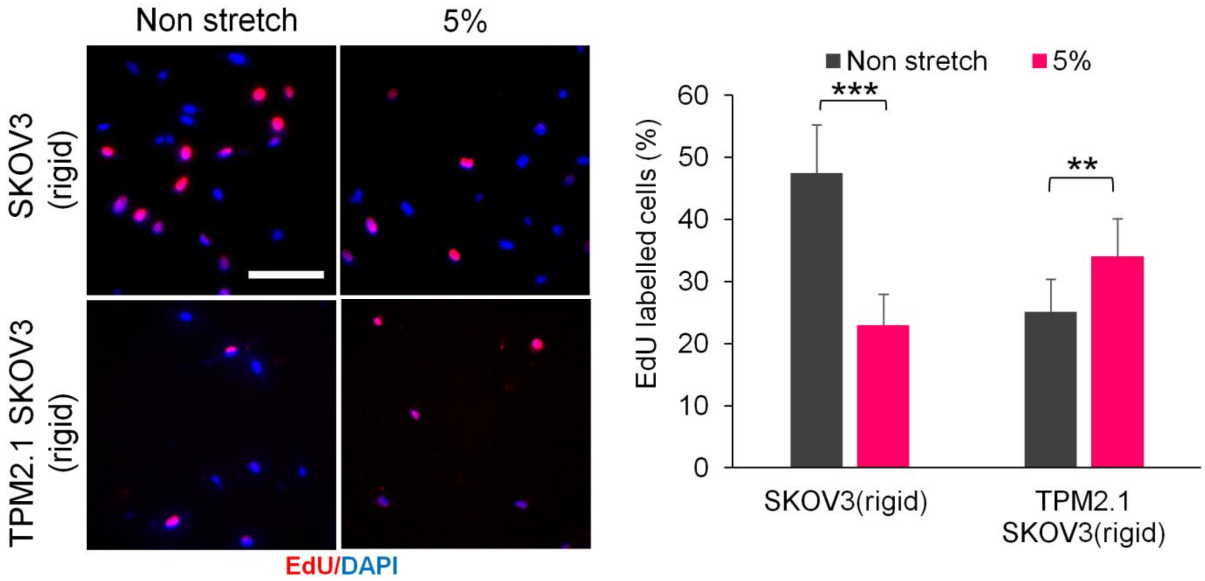


**Figure S5. Cyclic Stretch Reduces SKOV3 Cancer Cell Growth but TPM2.1 Expression Promotes Cell Growth**

SKOV3 cancer cells showed decrease in the proliferation in response to cyclic stretch, while TPM2.1 expression enhanced cell proliferation with cyclic stretch, n>600 SKOV3 cells and n>250 TPM2.1 restored SKOV3 cells, scale bar: 100 µm, ***P*< 0.01, ****P*< 0.001.

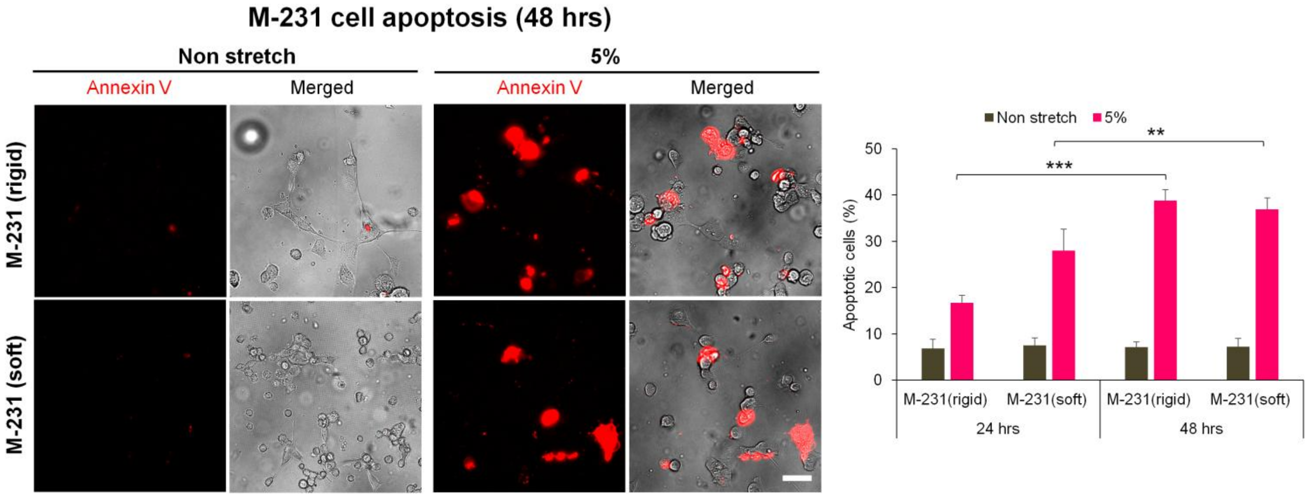


**Figure S6. Prolonged Cyclic Stretch Potentiates Cancer Cell Apoptosis**

Cyclic stretching of M-231 cancer cells for 48 hrs further potentiated cell apoptosis on rigid (37%) and soft (39%) surfaces compared to 24 hrs of cyclic stretching (16% on rigid) and (28% on soft surface). n>1000 cells for 24 hrs assay, cells were from four independent experiments and n>250 cells for 48 hrs assay, cells were from two independent experiments, scale bar: 50 µm, ***P*< 0.01, ****P*< 0.001.

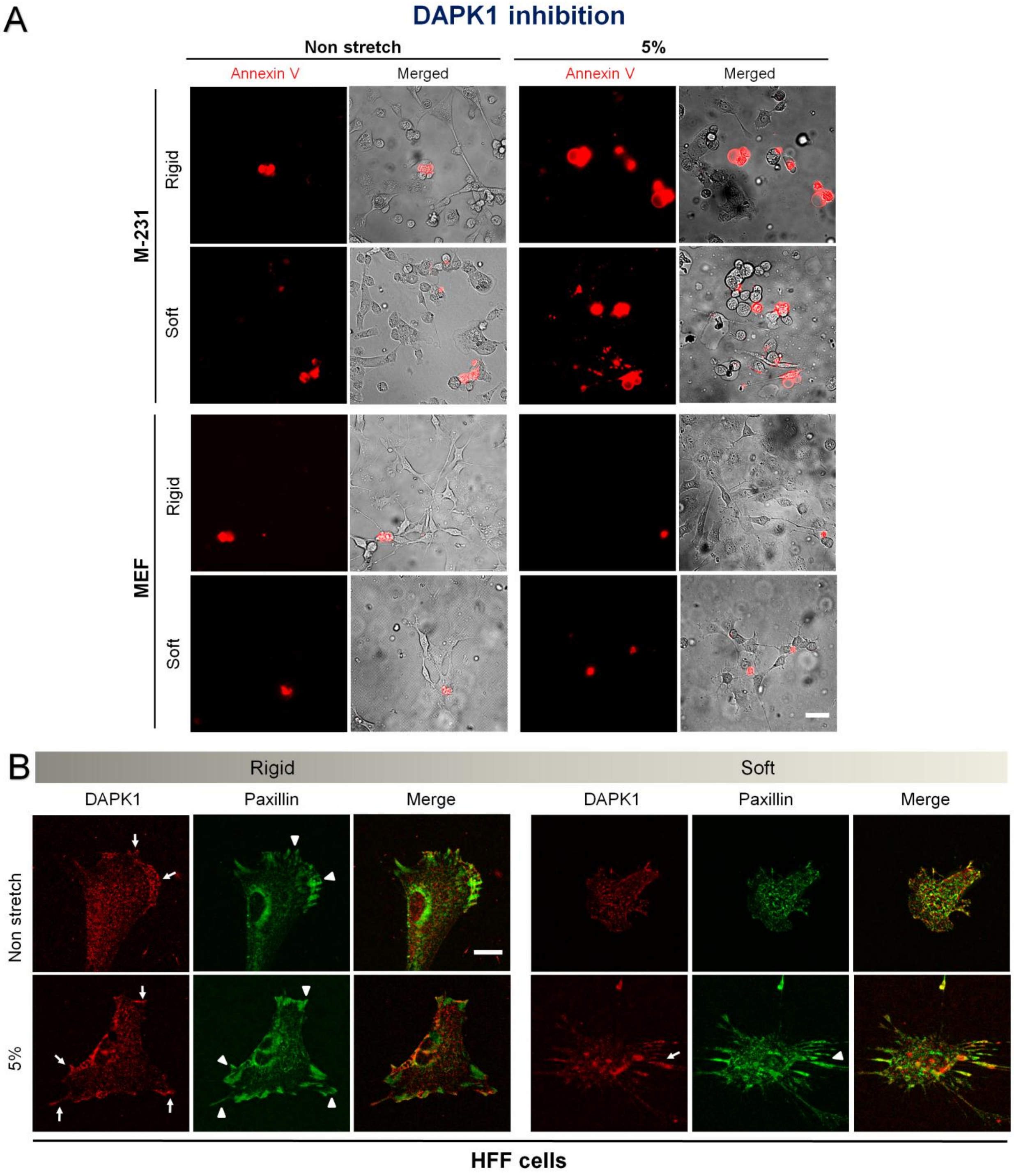
A
DAPK1 inhibition
Non stretch
5%
Annexin V
Merged
Annexin V
Merged
M-231
Rigid
Soft
MEF
Rigid
Soft
B
Rigid
Soft
DAPK1
Paxillin
Merge
DAPK1
Paxillin
Merge
Non stretch
5%
HFF cells

**Figure S7. DAPK1 inhibition Reduces Apoptosis in Normal Cell Grown on Soft Surface but not in Cancer Cell**

**(A)** Representative images displaying the M-231 and MEF cell apoptosis in the presence of DAPK1 inhibitor. DAPK1 inhibition rescued MEF cell apoptosis on non stretched soft surface but not in the M-231 cancer cell, scale bar: 50 µm. **(B)** Representative images showing colocalization of DAPK1 and paxillin not only in stretched and non stretched HFFs on rigid surfaces but also in the cells on stretched soft surface. However, such colocalization was absent in the cells on non-stretched soft surface, indicating that DAPK1 is activated in those cells. White arrow indicates DAPK1 localization and arrow head denotes paxillin position at the same location. Scale bar: 20 µm

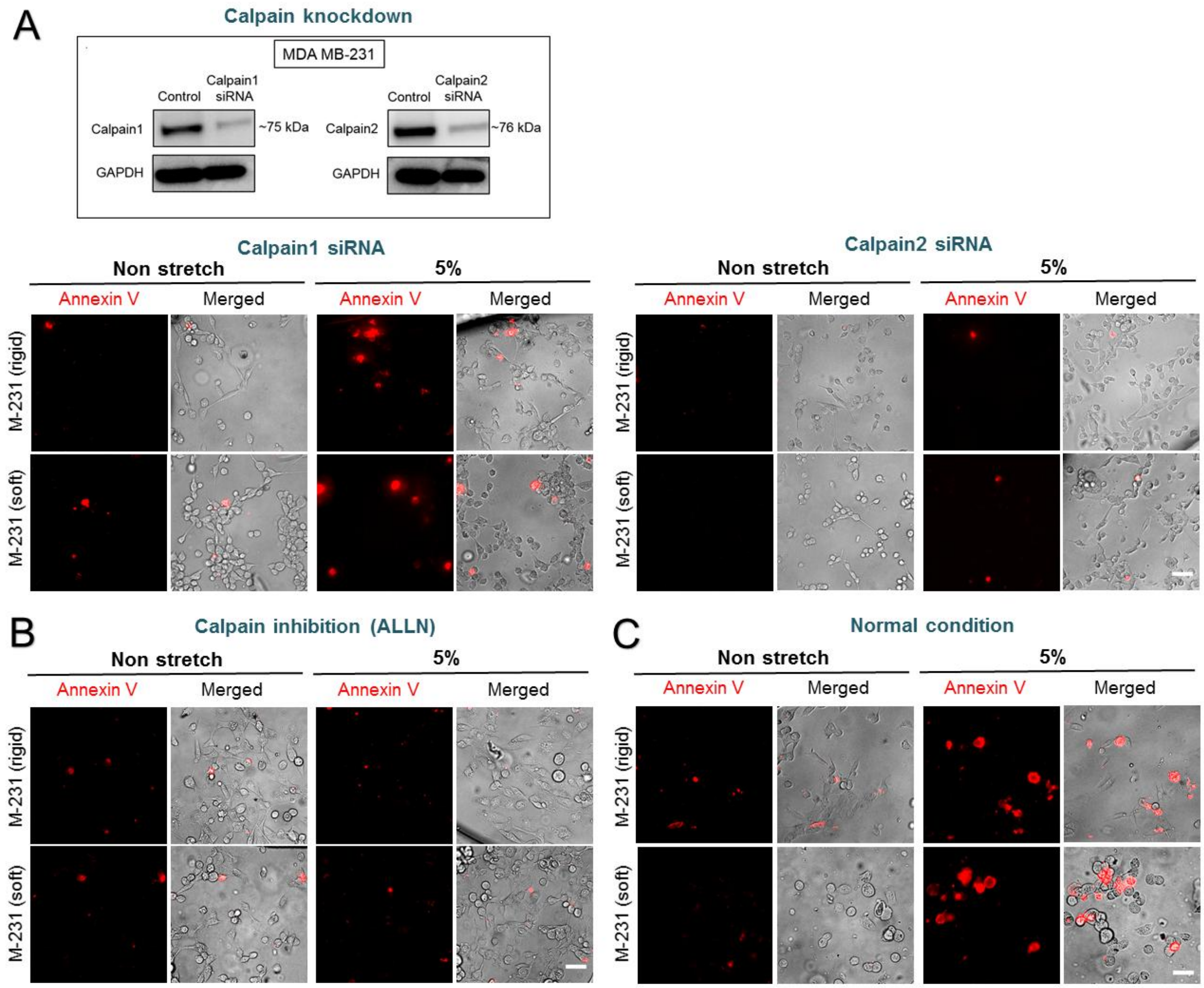


**Figure S8. Calpain Inhibition decreased Apoptosis in Cyclically Stretched M-231 Cancer Cells**

**(A)** Western blot data illustrating specific knockdown of Calpain-1 and -2 using siRNAs. Representative images depicting a substantial reduction in the apoptosis in calpain-1 and -2 knockdown cancer cells respectively, n>1000 cells. **(B)** Representative images showing similar decrease in the apoptosis in ALLN (calpain-1 and -2 inhibitor) treated cancer cells, n>500 cells. **(C)** Representative images of untreated cancer cells (as a control) illustrating the stretch-induced apoptosis in those untreated cancer cells, n>500 cells. All scale bars: 50 μm

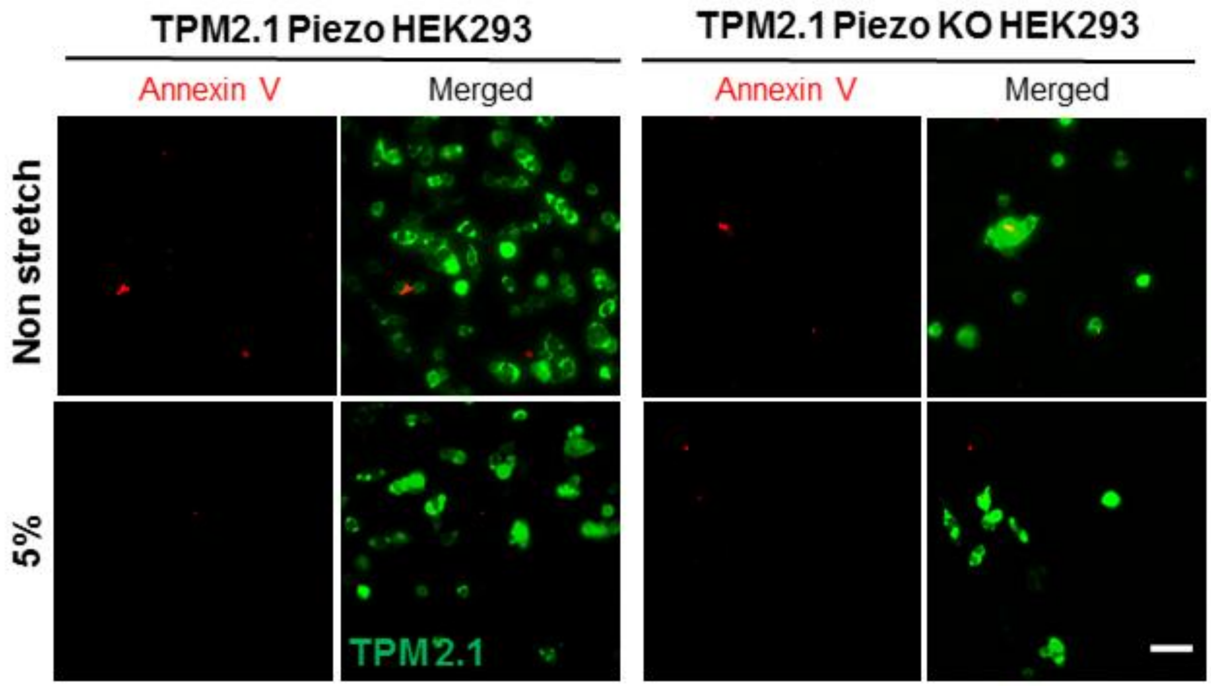


**Figure S9. TPM2.1 Restoration in Transformed HEK293 Cells Down-regulates Apoptosis**
Representative images showing a reduction in the apoptosis in TPM2.1 restored HEK293 cells. Scale bar: 50 µm.
